# Whole-embryo Spatial Transcriptomics at Subcellular Resolution from Gastrulation to Organogenesis

**DOI:** 10.1101/2024.08.27.609868

**Authors:** Yinan Wan, Jakob El Kholtei, Ignatius Jenie, Mariona Colomer-Rosell, Jialin Liu, Joaquin Navajas Acedo, Lucia Y. Du, Mireia Codina-Tobias, Mengfan Wang, Ahilya Sawh, Edward Lin, Tzy-Harn Chuang, Susan E. Mango, Guoqiang Yu, Bogdan Bintu, Alexander F. Schier

**Author notes:** These authors contributed equally to this work (Jakob El Kholtei, Ignatius Jenie). Corresponding authors. (Y.W.); (B.B.); (A.F.S.).

## Abstract

Spatiotemporal patterns of gene expression underlie embryogenesis. Despite progress in single-cell genomics, mapping these patterns across whole embryos with comprehensive gene coverage and at high resolution has remained elusive. Here, we introduce a <u>w</u>hole-<u>e</u>mbryo imaging platform using <u>m</u>ultiplexed <u>e</u>rror-robust fluorescent in-<u>s</u>itu <u>h</u>ybridization (weMERFISH). We quantified the expression of 495 genes in whole-mount zebrafish embryos at subcellular resolution. Integration with single-cell multiomics data generated an atlas detailing the expression of 25,872 genes and the accessibility of 294,954 chromatin regions, explorable with an online interface MERFISHEYES (beta version). We found that temporal gene expression aligns with cellular maturation and morphogenetic movements, diverse expression patterns correspond to composites of tissue-specific accessible elements, and changes in gene expression generate sharp boundaries during gastrulation. These results establish a novel approach for whole-organism spatial transcriptomics, provide a comprehensive spatially resolved atlas of gene expression and chromatin accessibility, and reveal the diversity, precision and emergence of embryonic patterns.

## Introduction

Understanding the spatiotemporal patterns of gene expression is crucial for deciphering the principles that underlie animal development and disease states. Historically, the expression of individual genes has been detected by ever evolving RNA visualization techniques (*1–4*), resulting in resource databases consisting of large collections of *in situ* stainings across embryos of different model organisms (e.g., Zebrafish Informational Network (*5, 6*) and Berkeley Drosophila Genome Project (*7–9*)). Over the past decade significant advances in high-throughput spatial omics techniques have enabled simultaneous measurements of many genes in the same embryos to identify co-expression patterns (*10*). For example, sequencing-based spatial transcriptomics methods (e.g., Tomo-seq (*11*) and Stereo-seq (*12*)) have mapped the transcriptome across 2D sections of developing zebrafish. Whole mouse embryos were reconstructed with a method called Slide-seq (*13*) by realigning many consecutive serial sections. While in principle covering the entire transcriptome, these methods only cover the more abundant genes due to low capture efficiency, and provide low spatial resolution, typically not at the single-cell level. Alternative image-based spatial transcriptomics methods such as MERFISH (*14, 15*) or seqFISH (*16–18*) provide high subcellular resolution for 2D tissue sections and high detection efficiency, capturing >50% of transcripts, albeit with a more limited targeting ability of a few hundred genes. However, these methods are not currently compatible with imaging entire embryos hundreds of microns in size. Recent computational methods (*19*) have improved the limited targeting constraint of imaging-based spatial transcriptomics methods by mapping single-cell transcriptomes measured via scRNA-seq onto reference spatial transcriptomics atlases, thus allowing a more comprehensive view of gene expression patterns. Similar approaches were used for zebrafish and *Drosophila* embryos (*20–22*), but the low resolution of the reference atlases and limited numbers of landmark genes restricted transcriptome mapping to low spatial resolution and embryos with simple geometry. Thus, constructing detailed atlases of single mRNA molecules with subcellular resolution across entire 3D embryos remains a challenge and has hindered the systematic study of fundamental questions in developmental gene expression: What is the full diversity of gene expression patterns? How precisely do these patterns define positional information? How are gene expression patterns linked to the differential accessibility of genomic elements? How do these patterns evolve over time?

### weMERFISH detects the mRNAs of hundreds of genes at subcellular resolution in whole zebrafish embryos

To address the limitations of existing spatial-omics methods, we developed a novel imaging platform to profile spatial gene expression in whole-mount embryos from gastrula to organogenesis stages, using zebrafish as a model system (Fig.1A). Four major adaptations were implemented compared to conventional MERFISH methods (*14, 15*) to achieve high-resolution single-molecule imaging across the whole embryo (Fig. 1B): (1) Probe anchoring: the primary DNA probes hybridizing to the RNA were synthesized with an acrydite-modification, allowing for their covalent incorporation into protective polyacrylamide gels casted on top of the embryo samples. This approach enables long-term sample stability (>1 month) regardless of mRNA degradation and complete removal of fluorescent probes using 80-100% formamide, allowing repeated imaging over more than 60 rounds. (2) Codebook flexibility: instead of designing the MERFISH combinatorial code in the primary probes, we designed the primary probe readout sequences in a gene-specific manner and used a mixture of linker probes to combinatorially encode and image the gene library via MERFISH. This approach allows convenient switching between imaging a single gene with single-molecule FISH versus different combinatorial strategies for MERFISH, allowing for facile adaptation and optimization. (3) Confocal imaging: we used spinning confocal microscopy that eliminates out-of-focus light in deep tissues. (4) Signal amplification: to address the signal loss in confocal imaging, we implemented a two-level branching signal amplification (*23, 24*) that increased the signal by three-fold (Fig. S1).

**Fig. 1.**
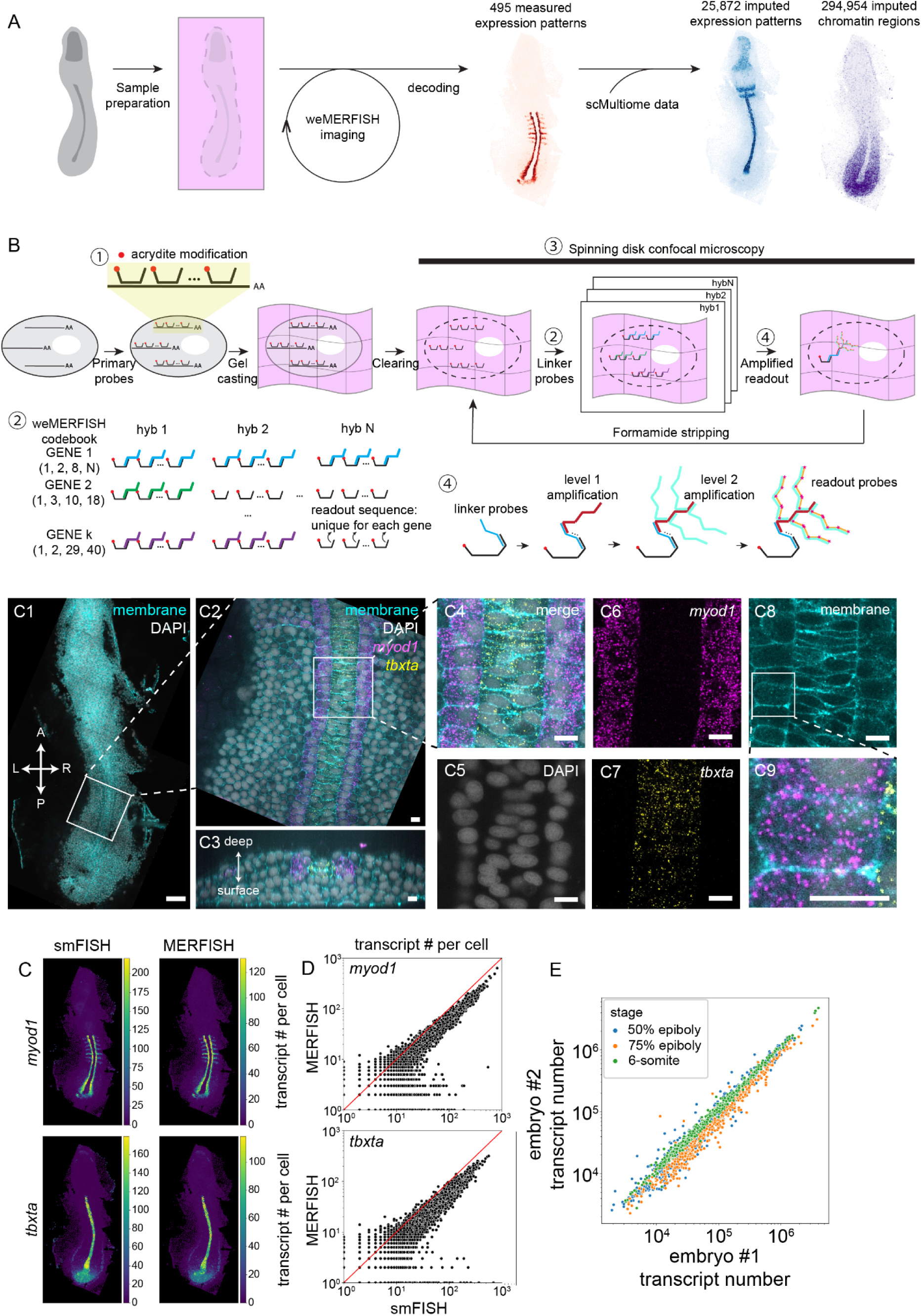
weMERFISH enables imaging of whole zebrafish embryos for 495 genes. **(A)** Sample preparation, imaging and data processing pipeline: The zebrafish embryo is embedded in a polyacrylamide gel, which become transparent after clearing of proteins and lipids; weMERFISH imaging was performed and single transcript locations were decoded from the imaging data; by incorporating single-cell multiome data, n=25,872 gene expression patterns and n=294, 954 chromatin accessible regions can be mapped to the weMERFISH atlas **(B)** Sample preparation and imaging pipeline for weMERFISH: (1) Zebrafish embryos are embedded in polyacrylamide gels to crosslink acrydite-modified primary probes. (2) A MERFISH combinatorial design is implemented by mixing gene-specific linker probes for each round of hybridization. (3) The embryos are imaged with a spinning disk confocal microscope upon (4) a branching tree signal amplification strategy. **(C)** Representative images of two genes (*myod1* (magenta) and *tbxta* (yellow)) visualized by the serial single-molecule FISH component of weMERFISH: (C1) Overview of an unraveled 6-somite stage zebrafish embryo. Membrane staining is shown in cyan, and DAPI (nuclear stain) in gray. (C2) A zoomed-in view of a single z-plane. (C3) A zoomed in cross-sectional view marking the bottom and deeper tissue layers. (C4-8): Progressive zoom-in views of the embryo in B1 to subcellular resolution. **(D)** *myod1* and *tbxta* expression patterns of transcripts/cell quantified with the smFISH and MERFISH modalities of weMERFISH. **(E)** Correlation of transcript numbers per cell in smFISH versus MERFISH. Correlation coefficients for *myod1*: R² = 0.924, slope = 0.597, intercept = -0.059 and *tbxta*: R² = 0.890, slope = 0.552, intercept = 0.663. **(F)** Correlation of total number of transcripts per gene between replicate embryos for the three stages imaged. Total correlation coefficient: R = 0.996. Scale bars: 100 µm (C1-C3); 10 µm (C4-C9).

With the probe library we targeted 495 genes that were chosen based on their variable expression across cell types and/or time points in single cell/nucleus sequencing datasets (*25, 26*) or prior embryogenesis literature (Supplementary DataS1). The gene set included 217 transcription factors, 191 morphogenesis-related genes, and 59 markers associated with mature cell types or cell cycle progression (see Methods, Fig. S2). 471 genes were imaged in an 80-bit MERFISH codebook in 40 hybridization rounds; to avoid molecular crowding, 24 highly-expressed genes were imaged sequentially in 12 hybridization rounds.

Zebrafish embryos at 3 different stages (onset of gastrulation (50% epiboly), mid-gastrulation (75% epiboly), and early organogenesis (6-somite stage)) were unraveled from their yolks and then imaged (Fig. 1C). Individual RNA molecules were identified and assigned to cells segmented based on nuclear and membrane staining using machine learning-based methods (*27*)(Methods). The MERFISH results were highly correlated with the same genes imaged sequentially using single-molecule fluorescent in-situ hybridization (smFISH), with a recovery rate of 50-90% and false positive rate of <1 transcript per gene per cell (Fig. 1D-E and Fig. S3A). Two embryos were imaged for each stage, encompassing a total of 103,646 cells. The reproducibility between replicate embryos was very high (log-log correlation R=0.996, Fig. 1F), and the result showed good correlation with stage-matched bulk-RNA-seq data (log-log correlation R=0.66, Fig. S3B) (*28*). These results establish weMERFISH as an imaging platform that captures whole-embryo spatial transcriptomes at subcellular resolution.

### weMERFISH defines diverse gene expression patterns and cell types

To explore the weMERFISH expression patterns of the 495 genes, transcripts were visualized after the unraveled embryos were reassembled in 3D (Fig. S4). The weMERFISH data displayed remarkable expression diversity across the entire embryo (red panels in Fig. 2). Specific patterns of gene expression marked all major tissue types and regions (e.g. *rx3*, *mab21l2* and *six7* in the optic cup; *mef2d*, *myod1* and *prdm1a* in adaxial cells; *etv2* and *gfiaa* in the lateral plate mesoderm; *foxd3* and *sox10* in the neural crest). weMERFISH gene expression patterns displayed strong congruence to published data. For example, *tbxta*, *bambia* and *sox32* had identical or very similar expression patterns in weMERFISH embryos compared to published data (*5, 29–32*)(Fig. S5A). In some cases, weMERFISH helped further refine published expression patterns, as colorimetric *in situ* data can obstruct the detection of gene expression in overlaying domains. For example, we identified genes expressed in both the mesendoderm (hypoblast) and the overlaying ectoderm (epiblast) during mid-gastrulation with a class of genes having a similar expression pattern between the two layers (e.g., *her5*, *irx1b*, *irx7*, *mych*, *meis3* and *admp*) and a class of genes with divergent expression pattern (e.g. *otx1, otx2a*, *ras11b*, *zic2a*, *zic3* and *tcf7l1a*) (Fig. S5B). These results demonstrate the power of weMERFISH in detecting the precise expression of hundreds of genes in whole embryos.

**Fig. 2.**
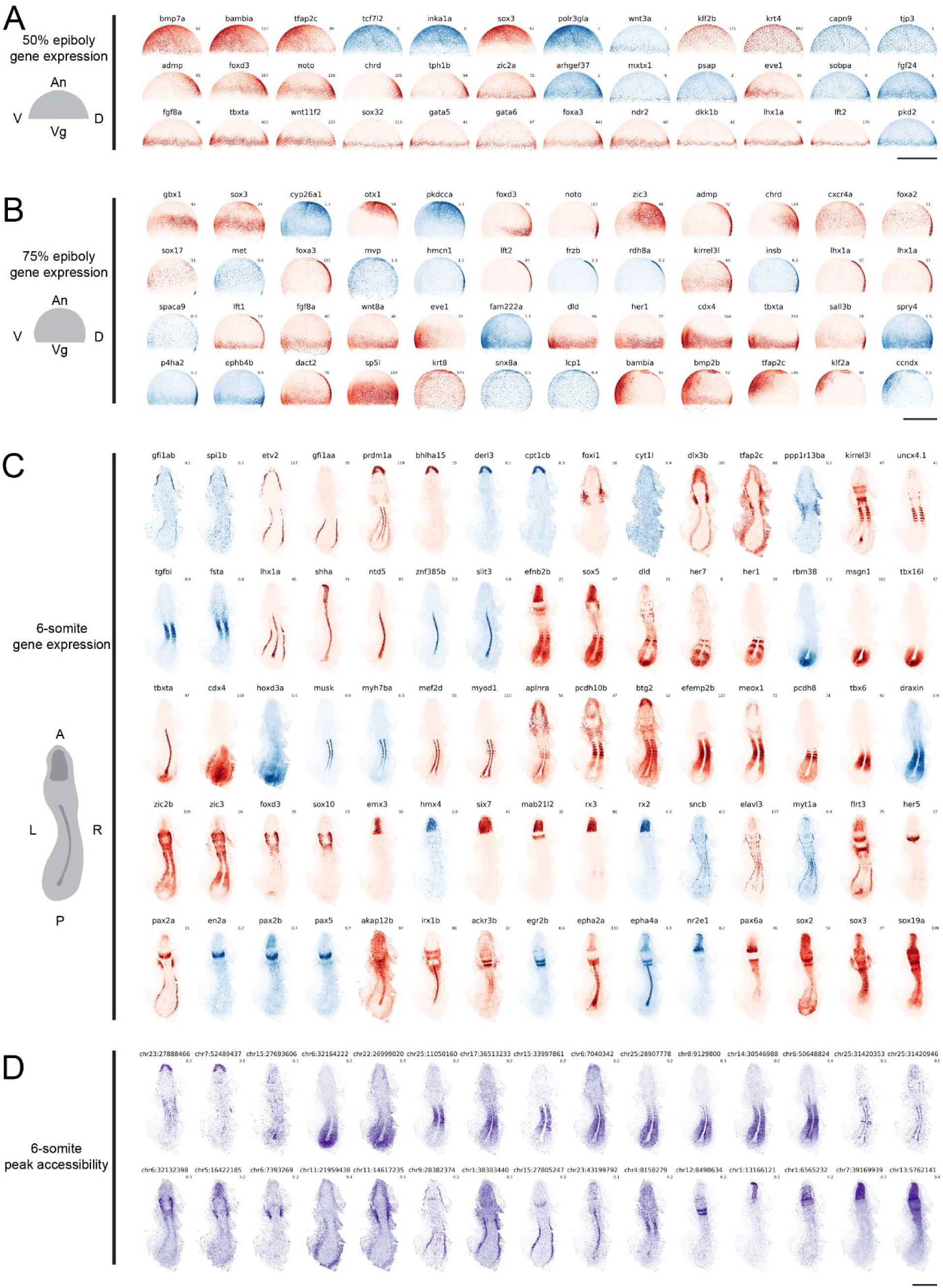
Diverse gene expression and chromatin accessibility patterns in early embryos. This figure showcases the spatial distribution of gene expression and chromatin accessibility during different stages of early zebrafish embryogenesis. Gene expression patterns measured by weMERFISH are shown in red, imputed gene expression patterns from scMultiome sequencing data (*26*) in blue, and imputed chromatin accessibility patterns in purple. **(A-B)** 50% and 75% epiboly gene expression: The expression patterns of various genes are visualized at the 50% and 75% epiboly stages from the lateral view. The schematic on the left illustrates the dorsal (D), ventral (V), anterior (An), and vegetal (Vg) regions of the embryo, corresponding to the respective expression patterns shown. Genes such as *bmp7a*, *sox3*, *wnt3a*, *foxa2*, and others are highlighted, illustrating distinct regional expression profiles within the embryo. **(C)** 6-somite stage gene expression: The expression patterns of genes are visualized at the 6-somite stage. The schematic on the left indicates the anterior (A), posterior (P), left (L), and right (R) sides of the embryo. Genes such as *myod1*, *tbxta*, *sox10*, *six7*, *meox1*, and others display distinct expression patterns across different regions of the developing embryo. **(D)** 6-somite stage peak chromatin accessibility: The chromatin accessibility patterns of genomic loci neighboring key developmental genes such as *lhx1a* (chr15:27805247), *myod1* (chr25:31420946) and *mespbb* (chr25:11050160) are shown at the 6-somite stage. Chromosomal location indicate peak start, and the peak width is 500 bp. Scale bars: 500 μm

weMERFISH also opens the possibility to identify patterns that might not be easily detectable via traditional colorimetric *in situs*. We performed differential gene expression analysis between (1) spatially distinct regions at the 6-somite stage and (2) along the dorsal-ventral and animal-marginal axes of the early gastrula. We discovered two zones that have not been described previously. At the 6-somite stage, we identified a zone of specific gene expression in the anterior portion of the notochord posterior tip, between the more mature differentiating notochord anteriorly and the proliferative notochord progenitors posteriorly. Genes downregulated in this zone included the G1/S transition factors *mcm2, mcm3, mcm4, mcm7* and *uhrf1*, while *cdkn1ca*, a negative regulator of cell cycle, is up regulated (Fig. S6A). At the onset of gastrulation, we identified a set of 83 genes that are expressed at lower levels at the embryonic margin (Fig. S6B), most of which are zygotically expressed (Fig. S6C). GO enrichment analysis indicates that this gene set includes factors implicated in mitotic DNA replication initiation, chromatin organization and cell cycle (Fig. S6D). The gastrula margin has been considered a zone where mesendodermal genes are activated, whereas the large-scale gene repression found here has not been reported. These observations illustrate the potential of weMERFISH in discovering new global embryonic expression domains.

We next explored if the subcellular resolution of weMERFISH can discover RNA localization patterns within individual cells. We measured the average distance of each transcript species to the cell membrane. Strikingly, we found that subcellular localization is a gene-specific feature, with highly reproducible patterns between embryos (Fig. S7A). For example, in the deep cells of the gastrula (50% epiboly stage), transcripts close to the membrane include *akap12b, kank1a, lima1a,* where *akap12b* is localized to cell membranes in all cells during gastrulation, and to notochord and adaxial cells at the 6-somite stage (*33*)(Fig. S7B). Transcripts far away from the membrane include *her1, kmt2bb, lamb1a, cdx4, add3b, aspm, gbx1*, and *efnb2b* (Fig. S7A). Notably, transcript localization patterns can also dynamically change across stages. For example, Znf185 transcripts are diffusely expressed in the enveloping layer (EVL) cells at the onset of gastrulation (50% epiboly stage), but localize to the EVL cell junctions at mid-gastrulation (75% epiboly stage) (Fig. S7C). These results highlight the potential of weMERFISH in the systematic discovery of localized RNAs.

Based on the mRNA molecules imaged in each cell, we clustered the cells into different types/states analogously to single cell sequencing methods (*34*) (Fig. 3). Almost all of these cell states formed well defined spatial domains that were highly stereotypical across embryos (Fig. S8). At 50% epiboly, the clusters occupy positions at the dorsal margin, ventrolateral margin, and animal pole, and extraembryonic clusters represent the enveloping layer (EVL) and yolk syncytial layer (YSL) (Fig. 3A and Fig. S8A).

**Fig. 3.**
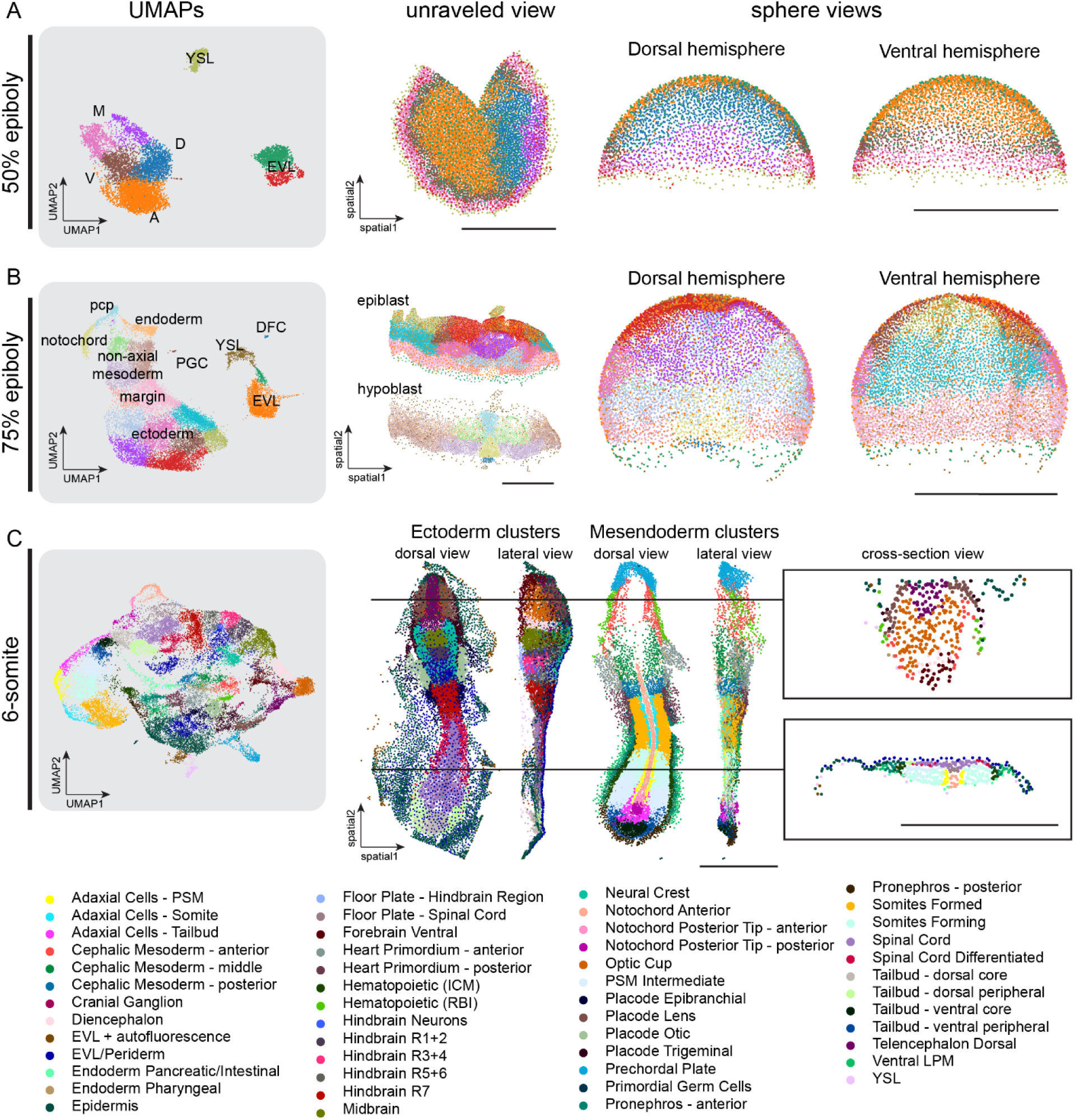
Clustering analysis of weMERFISH data across different zebrafish developmental stages. This figure illustrates the clustering of single-cell expression data obtained through weMERFISH at different stages of zebrafish embryogenesis, in transcriptional space using UMAP (Uniform Manifold Approximation and Projection) visualizations, and in physical space in the unraveled views, and sphere views of the embryo. **(A)** 50% epiboly stage: UMAP: Clustering of cells at the 50% epiboly stage, representing distinct cell populations across the embryo. The clusters are color-coded, indicating different regions such as the dorsal (D), ventral (V), animal (A), and marginal (M) areas, along with the enveloping layer (EVL) and yolk syncytial layer (YSL). Unraveled View: A 2D projection of the embryo in native imaging space, where clusters are mapped to their spatial positions. Sphere View: 3D reconstructions of the embryo’s dorsal and ventral hemispheres. **(B)** 75% epiboly stage: UMAP: Clustering of cells at the 75% epiboly stage, with distinct clusters representing cell types like the yolk syncytial layer (YSL), enveloping layer (EVL), notochord, ectoderm, and non-axial mesoderm. Unraveled View: A 2D projection of the embryo in native imaging space, with clusters corresponding to different tissue layers (epiblast and hypoblast) shown in their spatial context. Sphere View: 3D views of the dorsal and ventral hemispheres. **(C)** 6-somite stage: UMAP: Clustering of cells at the 6-somite stage, showcasing the diversity of cell populations during this stage of early organogenesis. Each cluster is color-coded, and the name of the tissue type is listed below. The dorsal and lateral views of the embryo, with clusters colored as in the UMAP are shown. Additionally, two cross-sectional views at different anterior-posterior levels provide a detailed look at the spatial distribution of cell populations across the embryo. This figure details the spatial organization of cell clusters defined based on their single-cell transcriptomes during critical stages of zebrafish development. Scale bars: 500μm

Median difference between the cluster locations of the replicate embryos is 27 μm (Fig. S9A). At 75% epiboly, the embryo proper can be divided into 14 clusters including primordial germ cells, dorsal forerunner cells, YSL and EVL cells (Fig. 3B and Fig. S8B). Median difference between the cluster locations of the replicate embryos is 22 μm which is of similar dimension to a single cell (Fig. S9B). At 6-somite stage, the embryo can be partitioned into 51 cell types at distinct locations (Fig. 3C and Fig. S8C). The proportion of individual cell types differ between embryos on average by less than ∼3%, showing high stereotypy of the embryonic tissue composition (Fig. S8).

scRNA-seq data is widely used to cluster cells according to type and state (*12, 25, 35*). The definition of cell types and states in the spatial context allowed us to identify cell types missed in the published scRNA-seq clusters (*25, 35*) (Fig. 3). For example, adaxial cells, which were previously defined as a single cell population, formed three subclusters that locate at the level of formed somites, forming somites, and tailbud. Similarly, the tailbud, the notochord (Fig. S6A) and the cephalic mesoderm can all be separated into finer anatomically-defined weMERFISH clusters with differential gene expression patterns (Fig. 3C). Reverse mapping of weMERFISH subclusters to the UMAP of single-cell RNA-seq data (Fig. S10) allowed sub-clustering and annotation according to precise spatial information. These results show the power of weMERFISH in providing a comprehensive view of the precise spatial arrangement of the embryonic cell types and states.

### Integration of weMERFISH and single-cell multiome creates an atlas for gene expression and chromatin accessibility at genome-wide scale

Various efforts have been made in imputing non-measured genes from single-cell transcriptomics data to spatial locations in embryo data (*12, 18, 20, 21*). However, previous attempts have been hindered by low numbers of landmark genes and the lack of single-cell spatial resolution in reference maps. Our high-resolution large-scale dataset allowed the integration of weMERFISH data with single-cell multiomics (scMultiome) data consisting of single-nuclei RNA-seq and ATAC-seq (*26*). By learning a matrix denoting the probability of finding each snRNA-seq cell in each cell of the weMERFISH data using Tangram (*36*), cell-type labels were transferred from the snRNA-seq cells to the weMERFISH cells with high-fidelity (Fig. S11-12). This approach defined the spatial expression of 25,872 genes (Fig. 1A, blue panels in Fig. 2). Imputed gene expression patterns agreed with in-situ data (*5, 37*) and added expression domains that were not in the 495-gene probe set. For example, *cyp26a1* is consistently expressed in the embryonic margin and dorsal ectoderm at 75% epiboly, and *egr2b* in hindbrain rhombomeres 3 and 5 at 6-somite stage (Fig. S13).

The imputation allowed for exploring the expression pattern for each gene in both transcriptional space (UMAP) as well as in physical space. Certain genes, e.g. the anterior-posterior positioning of the Hox gene expression is readily apparent in weMERFISH embryos but not in transcriptional space (Fig. S14). By exploring this comprehensive atlas, we found hundreds of genes whose specific expression had not been described previously. For example, at 6-somite stage, genes *cacna2d1a*, *tnni2b.1* and *si:dkeyp-69b9.3* are expressed in adaxial cells, *zgc:194210* is expressed in the otic placode, *itm2cb* is expressed in the tailbud, and *si:ch211-214c7.4/prr36a* is expressed in the optic cup. These results demonstrate the power of combining weMERFISH with scRNA-seq data to define expression patterns at genome-wide scale.

The availability of scMultiome data, which simultaneously measures both RNA expression and ATAC accessibility in single cells, allowed us to impute the spatial location of 294,954 accessible chromatin regions into the 6-somite embryos imaged by weMERFISH using the RNA measurements as an intermediate (Fig. 1A). Imputed ATAC peak accessibility patterns agreed with previous enhancer activity reporter studies. For example, we identified specific enhancers in the *foxd3* (*38*) and *tbxta* (*39*) loci, and various enhancers reported in recent publications (*40, 41*) (Fig. S15 and Supplemental DataS3). We found thousands of peaks enriched in specific regions and cell types, including prechordal plate, tailbud, somite, neurons, neural crest, hematopoietic cells and various brain regions (purple in Fig. 2). These results demonstrate the power of combining weMERFISH with scMultiome data to define the spatial patterns of chromatin accessibility at genome-wide scale.

The integration of weMERFISH and scMultiome data establishes genome-wide maps for the spatial expression of 25,872 genes and the spatial location of 294,954 ATAC peaks. To support the exploration of this dataset and atlas, we created a preliminary browser interface called MERFISHEYES (<u>schier.merfisheyes.com</u>), in its beta version, to visualize the expression patterns of all annotated genes and the spatial location of accessible regions. Users can select a gene of interest and visualize in 3D its whole-embryo expression at three stages of development. In addition, users can interactively visualize the imputed accessibility of neighboring genomic regions surrounding the selected gene at the 6-somite stage through an interactive genome browser. MERFISHEYES will help users explore the 3D expression patterns and putative regulation of genes of interest.

### Cellular maturation and morphogenetic movements parallel gene expression dynamics

The weMERFISH atlas provides snapshots of spatial gene expression at three stages of embryogenesis. To offer a temporal component to this atlas, we used two inference approaches: (1) integrating the weMERFISH data with scRNA-seq for which pseudotime trajectories were previously computed (*25*) and (2) analysis of the weMERFISH data of the nuclear vs. cytoplasmic RNA distribution (called RNA-velocity (*42, 43*)) to infer the direction of transcriptional change.

Cells along the developmental pseudotime trajectories were mapped onto the three stages of the imaged embryos using Tangram (*36*). The spatial location of cells along the trajectories were captured: for example, the trajectories leading to the somite fates map to 1) non-axial mesoderm at blastula stage, then 2) the margin at mid-gastrula stage, 3) the tailbud, anteriorly progressing to 4) forming somites, and finally 5) mature somites at 6-somite stage (Fig. 4A). Similarly, the trajectories of the notochord map to axial mesoderm at early gastrula stage, then trace to the dorsal margin at mid-gastrula, and progress from the posterior tip to anterior notochord at the 6-somite stage (Fig. S16A). Notably, these maps show the location of cells that have the molecular potential to give rise to more specialized and differentiated cells, and thus resemble specification maps rather than fate maps (*44, 45*)

**Fig. 4.**
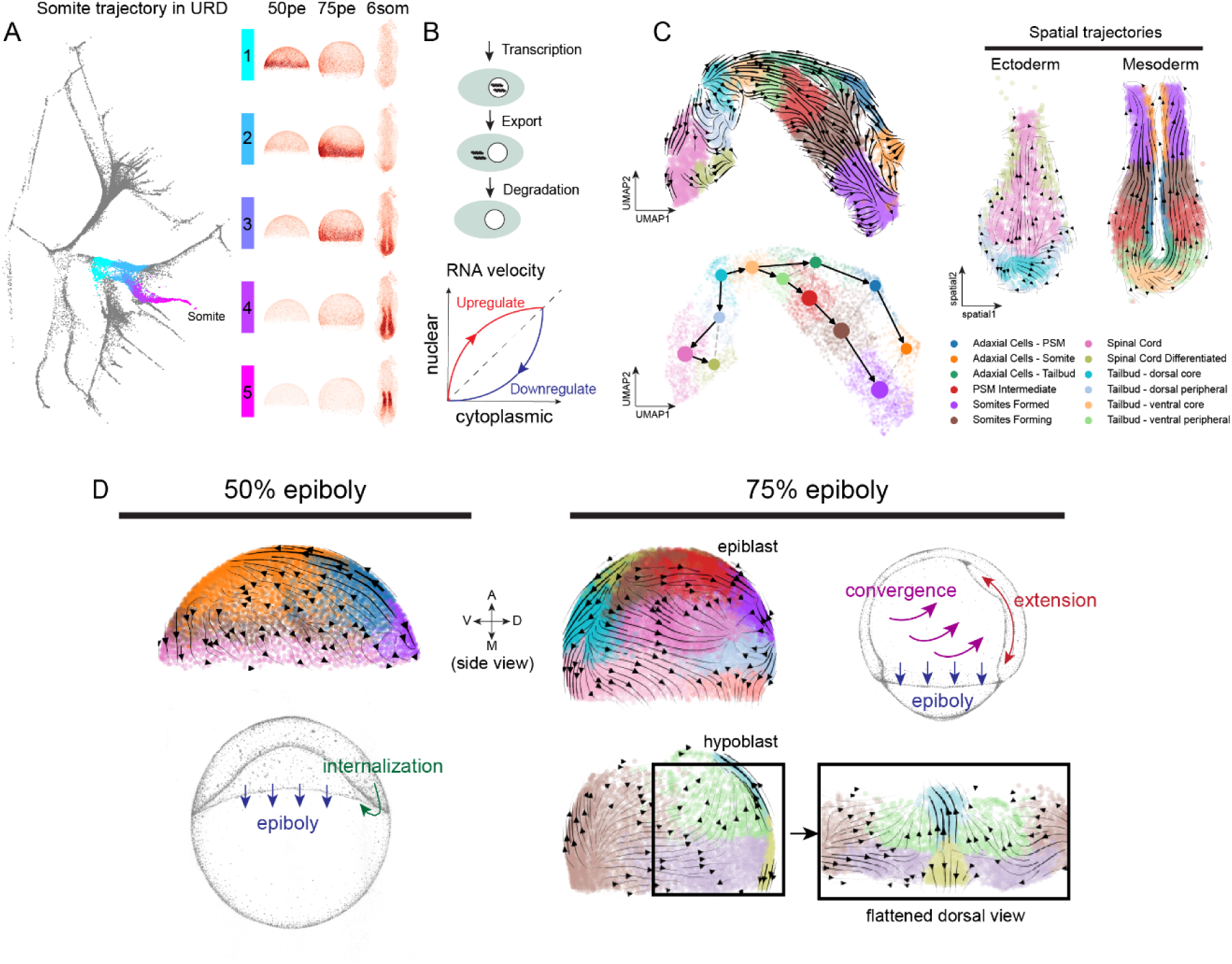
Developmental patterns inferred by pseudotime trajectory mapping and RNA-velocity analyses. **(A)** Mapping of pseudotime trajectories to physical space. Left: 5 segments along the pseudotime trajectory of somites are shown from single-cell sequencing data (*25*) in URD transcriptional space. Right: These stages are imputed to the spatial weMERFISH data across three developmental stages (50% epiboly, 75% epiboly and 6-somite stage). The segments trace the lineage progression from (1) the non-dorsal margin at 50% epiboly to (2) the paraxial mesoderm at 75% epiboly, to (3) tailbud (4) forming somite and (5) mature somite from posterior to anterior at 6-somite stage. **(B)** A **s**chematic illustration explaining the principle of the adapted RNA-velocity analysis (*42, 43*), which estimates the direction of transcriptional change by comparing the nuclear RNA abundance to cytosolic RNA abundance. The RNA velocity follows the prediction of gene expression dynamics, indicating whether transcripts are being upregulated (positive velocity and more nuclear accumulation) or downregulated (negative velocity and more cytosolic accumulation) **(C)** RNA velocity inferred for the tailbud trajectories using two computational methods (scVelo (*46*) - upper left panel) and (scVelo (*46*) combined with PAGA (*67*) - upper right panel). The inferred velocity streams are mapped to spatial locations within the embryo, estimating the directional flow of developmental processes within the ectoderm and mesoderm clusters. **(D)** Side-by-side comparison of RNA velocity map and morphogenetic movement at 50% and 75% epiboly stages. Upper-left: At 50% epiboly, RNA-velocity vectors match the internalization of cell during the epiboly process, with a directional movement towards the vegetal pole. Bottom-left: schematic of 50% epiboly morphogenic movement adapted from (*68*). Upper-right: At 75% epiboly, the epiblast and hypoblast RNA-velocity vectors match the convergence and extension movements critical for embryonic axis formation. A schematic of these processes is adapted for 75% epiboly from (*68*).

The subcellular resolution of weMERFISH allowed for quantifying nuclear vs. cytoplasmic transcript accumulation for each of the 495 genes imaged. As genes that initiate transcription have a bias towards nuclear accumulation, while genes terminating transcription have a bias for cytoplasmic accumulation, the nuclear/cytoplasmic transcript ratio allows for the reconstruction of the temporal direction of gene expression (*42, 43, 46*) (Fig. 4B). This approach identified developmental transitions in both transcriptional and physical space; e.g., from neuromesodermal progenitors in the dorsal tailbud to the differentiation of somites and spinal cord (Fig. 4C), and from posterior to anterior notochord (Fig. S16B-C). Strikingly, at the onset of and at mid gastrulation, velocity maps reflect cell morphogenetic patterns, with vectors aligning with the vegetal movement of marginal cells and the convergent extension of ventral-lateral cells (Fig. 4D). This observation suggests a connection between the temporal trend of gene expression dynamics and morphogenetic movements and can inform further investigations into the reciprocal regulation of gene expression and morphogenesis.

In summary, the development of weMERFISH and its integration with single-cell genomics datasets creates genome-wide whole-embryo views of spatiotemporal gene expression and chromatin accessibility. In the following sections we describe novel insights into the diversity, precision and emergence of patterned gene expression and regulation gained by the exploration of this data.

### Genes are often expressed in multiple related cell types and locations

The comprehensive mapping of spatial gene expression and accessible chromatin regions (Figs. 2 and 3) opens the opportunity to systematically analyze the tissue-specificity of developmental gene expression: How many genes define specific spatial domains and cell types? Are individual genes specific to one or multiple domains? Among the 3611 highly variable genes, analysis of 6-somite stage embryos identified 2275 genes that are enriched in the cells of the embryo proper (also called deep cells) (adjusted p-value<1e-50, log2 fold change>1). The highest number of specifically expressed genes is found in the prechordal plate, notochord and adaxial cells, potentially reflecting their status as early differentiating cells (Fig. 5A). The majority of genes (1316 out of 2275 differentially expressed genes) are expressed in two or more tissues or cell types. Notably, co-expression in two or more tissues is most frequently observed in cell types that share functional features or lineage relationships, such as notochord and prechordal plate (*47*); adaxial cells and somites; hindbrain rhombomeres; and forebrain regions (Fig. 5C). Additionally, we found that genomically proximate genes tend to be expressed in similar tissues (Fig. S17A), potentially reflecting the use of shared regulatory elements (*48*). This is well established for Hox genes (Fig. S14) but also applies to many other genomic loci. For example, a cluster of 15 genes including *ctslb*, *pank1b* and *slc16a12b* are co-localized in the genome and co-expressed in the hatching gland of the embryo. These results reveal the diversity of embryonic gene expression patterns and suggest that genes that share genomic locations have a higher likelihood to be expressed in multiple but related embryonic locations.

**Fig. 5.**
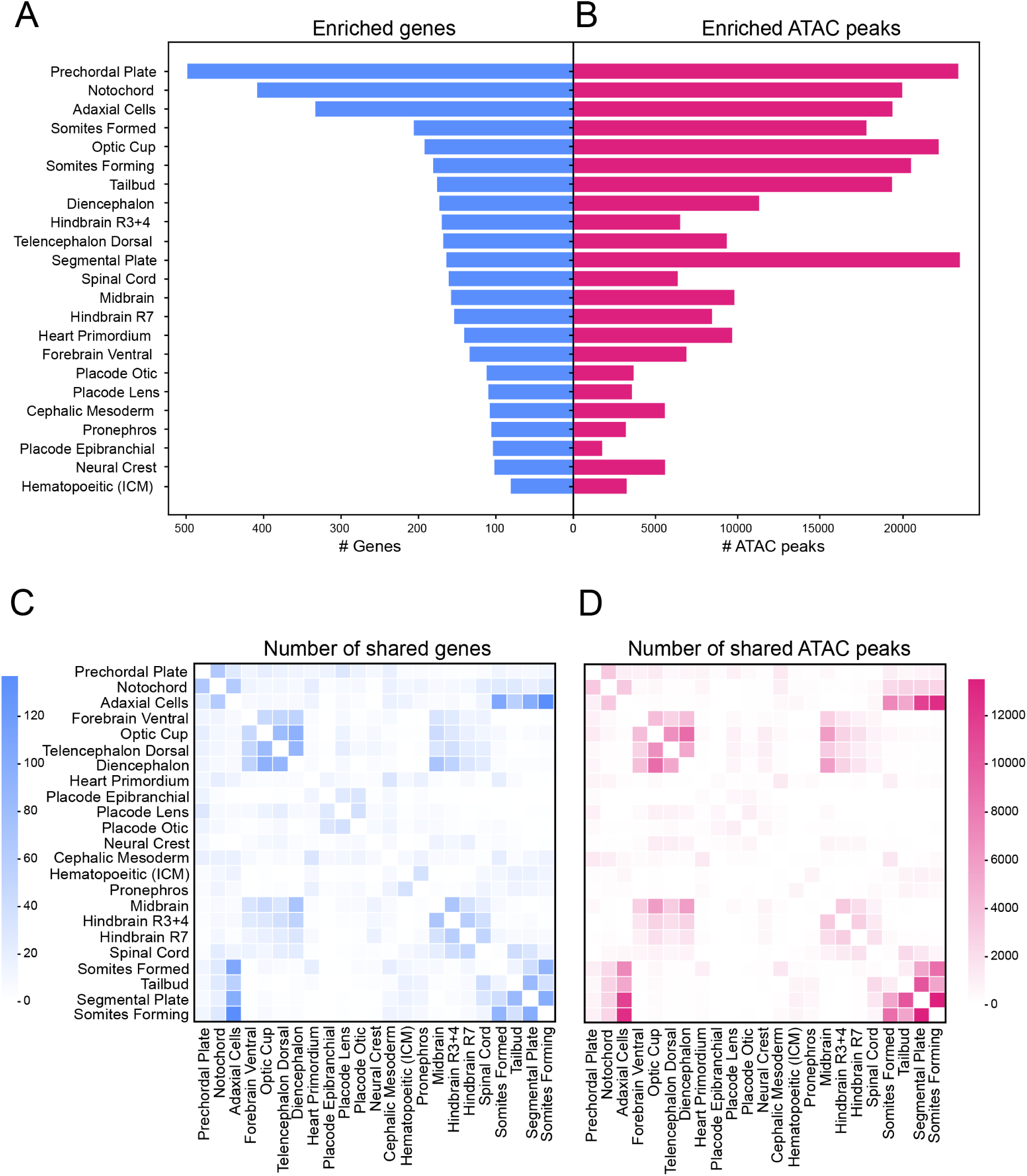
Tissue-specific gene expression and chromatin accessibility patterns in 6-somite stage embryo. (A) Number of genes enriched in specific tissues in the deep cell proper; (B) Number of ATAC peaks enriched in specific tissues in the deep cell proper; (C) Number of shared genes that are simultaneously expressed in the pair of tissues in the deep cell proper; (D) Number of shared ATAC peaks that are simultaneously accessible in the pair of tissues in the deep cell proper.

### Specific accessible chromatin regions combine to reflect complex expression patterns

The imputed atlas of accessible chromatin regions allows for correlating gene expression and putative cis-regulatory activity in space. Among the 294,954 ATAC peaks, 133,707 are enriched in the cells of the embryo proper (adjusted p-value<1e-50, log2 fold change>0.5) (Fig. 5B, D). As expected, genes enriched in specific tissues have genomic neighboring regions (<100 kb from the transcription start site (TSS)) with increased accessibility within the same tissue (Fig. S17B). For genes expressed in multiple tissues we asked if our data could distinguish two models of gene regulation (*49*). In one model, the accessibility of individual chromatin regions mirrors the transcription of neighboring genes: genes expressed broadly across tissues have corresponding putative regulatory elements with broad accessibility across the tissues. Alternatively, chromatin regions are accessible only in subsets of the regions a neighboring gene: putative regulatory elements combine to reflect the full expression pattern of a gene. We found that the majority of accessible regions are only enriched in a single tissue. For example, genes *hdlbpa* and *copz2* are expressed both in the notochord and prechordal plate, but 12/15 and 6/8 of their nearby ATAC peaks are accessible exclusively in notochord or prechordal plate (Fig. 6A). This single-tissue enrichment holds for all the 48 genes that are expressed in notochord and prechordal plate (Fig. 6B). Similarly, chromatin regions around dual-expressing genes in notochord and adaxial cells, or prechordal plate and adaxial cells also tend to be accessible only in one of the two tissues (Fig. S18). However, this trend does not apply to functionally and transcriptionally highly-related tissues, in which genes with shared expression in the two tissues tend to have genomic regions that are accessible in both tissues. For example, *palm1a* and *ghrhrb* are expressed in both adaxial cells and somites and are flanked by individual chromatin regions that are accessible in both tissues (Fig. 6C-D, Fig. S18). These results support the view that the specific expression profiles of individual genes across transcriptionally dissimilar tissues are composites generated by multiple tissue-specific regulatory elements, whereas genes expressed in related tissues are regulated by more broadly accessible elements.

**Fig. 6.**
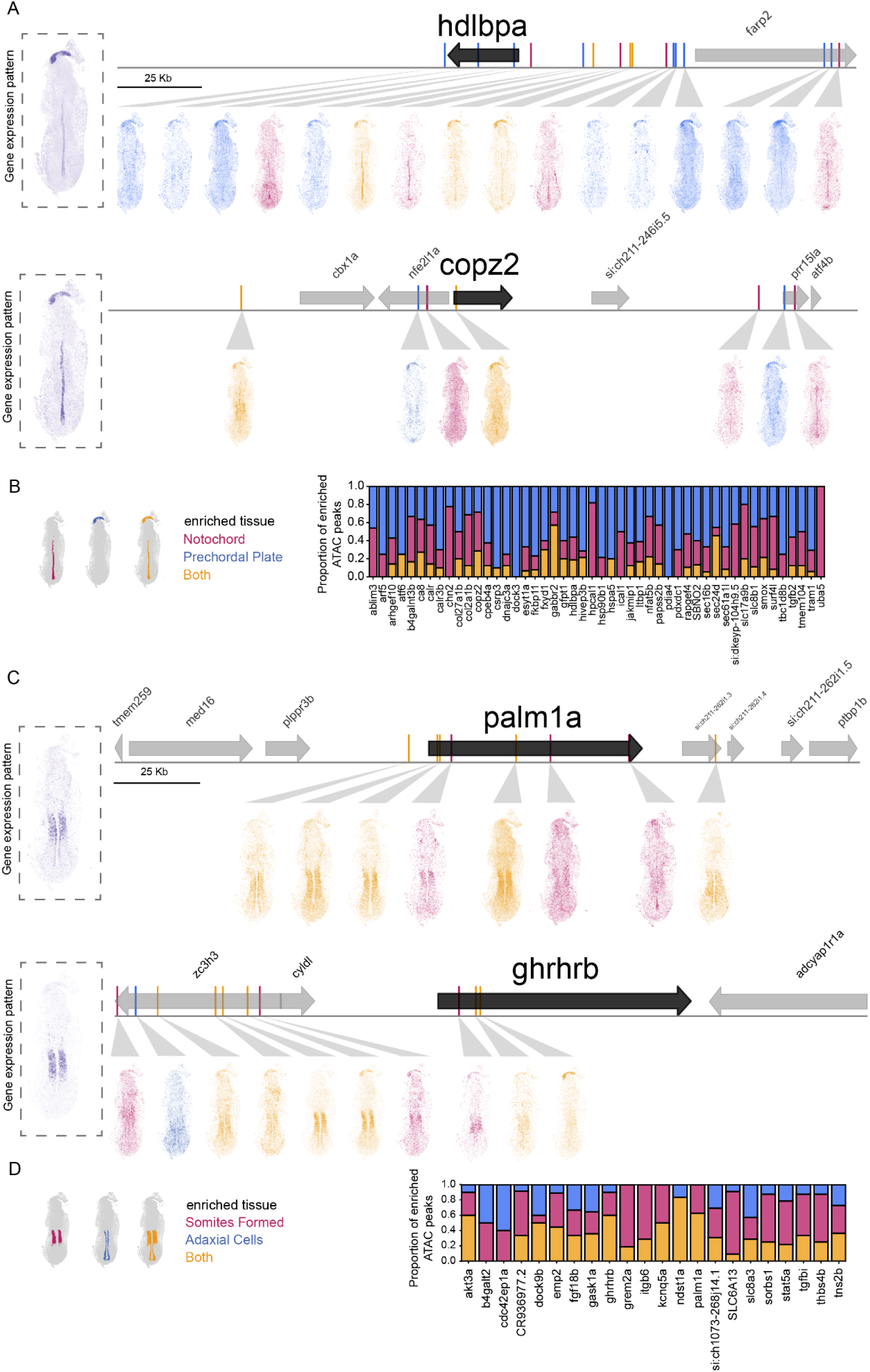
Regulatory logic of chromatin accessibility in different tissues. **(A-B)** Notochord and prechordal plate: (A) Two example genes (*hdlbpa* and *copz2*) with dual-tissue gene expression pattern (purple) are shown. The chromatin accessibility patterns (100 kilobases from the gene transcription start site (TSS)), showing mostly tissue-specific peaks in the notochord (pink) or prechordal plate (blue) with few shared between the two tissues (yellow). (B) Proportion of tissue-specific and shared accessible peaks neighboring all genes shared between the notochord and prechordal plate. **(C-D)** Formed somites and adaxial cells (C) Example genes (*palm1a* and *ghrhrb*) show different patterns, with more dual-tissue peaks for somites formed (pink) and adaxial cells (blue). (D) Same as (B) for somites formed and adaxial cells. Quantification reveals that overall tissue-specific peaks dominate divergent dual-tissue patterns, suggestive of independent regulatory control of genes share across divergent tissues.

### The formation of sharp boundaries is associated with gene expression changes

The availability of hundreds of gene expression patterns in the same embryo provides the opportunity to analyze two important developmental concepts: the extent of positional information and the emergence of precise gene expression territories. How distinct are neighboring cells across spatial domains? How do these distinctions arise during embryogenesis? To address the first question, we analyzed gene expression patterns along the anterior-posterior axis in the brain in the 6-somite embryos. We found that patterned expression is remarkably diverse and precise (Fig. S19), as even directly neighboring cells can be distinguished through a combination of genes. Interestingly, we found two broad categories of genes: those that mark a tissue with clear borders with their neighbors, and those that are expressed in a staggered pattern within a region. For example, genes such as *lhx2b*, *lhx9*, *six7* and *rx2* mark the entire optic cup, whereas combinations of staggered genes such as *barhl2*, *otx1*, *her9*, *en2b* and *pax5* are positioned continuously within the forebrain and midbrain. These results reveal remarkably high positional information in the developing embryo.

To study the emergence of gene expression territories, we characterized the differences in gene expression patterns during mid-gastrulation (75% epiboly stage) by measuring the heterogeneity of transcription within 50 μm of each cell (Fig. 7A, see methods). This approach revealed that notochord progenitors and somitic mesoderm progenitors formed the sharpest gene expression boundary (Fig. 7B, Fig. S20A). Across this border many genes exhibit sharp transitions from low to high expression within 30 μm (Fig. 7C, Fig S20B-C). In contrast, gene expression differences between other cell types were less uniformly sharp (e.g., the notochord-prechordal plate border, the dorsal-lateral posterior ectoderm border). Individual genes display sharp transitions, but the positions of their boundaries are staggered within the border region (Fig, 7C, Fig S20B-C).

**Fig. 7.**
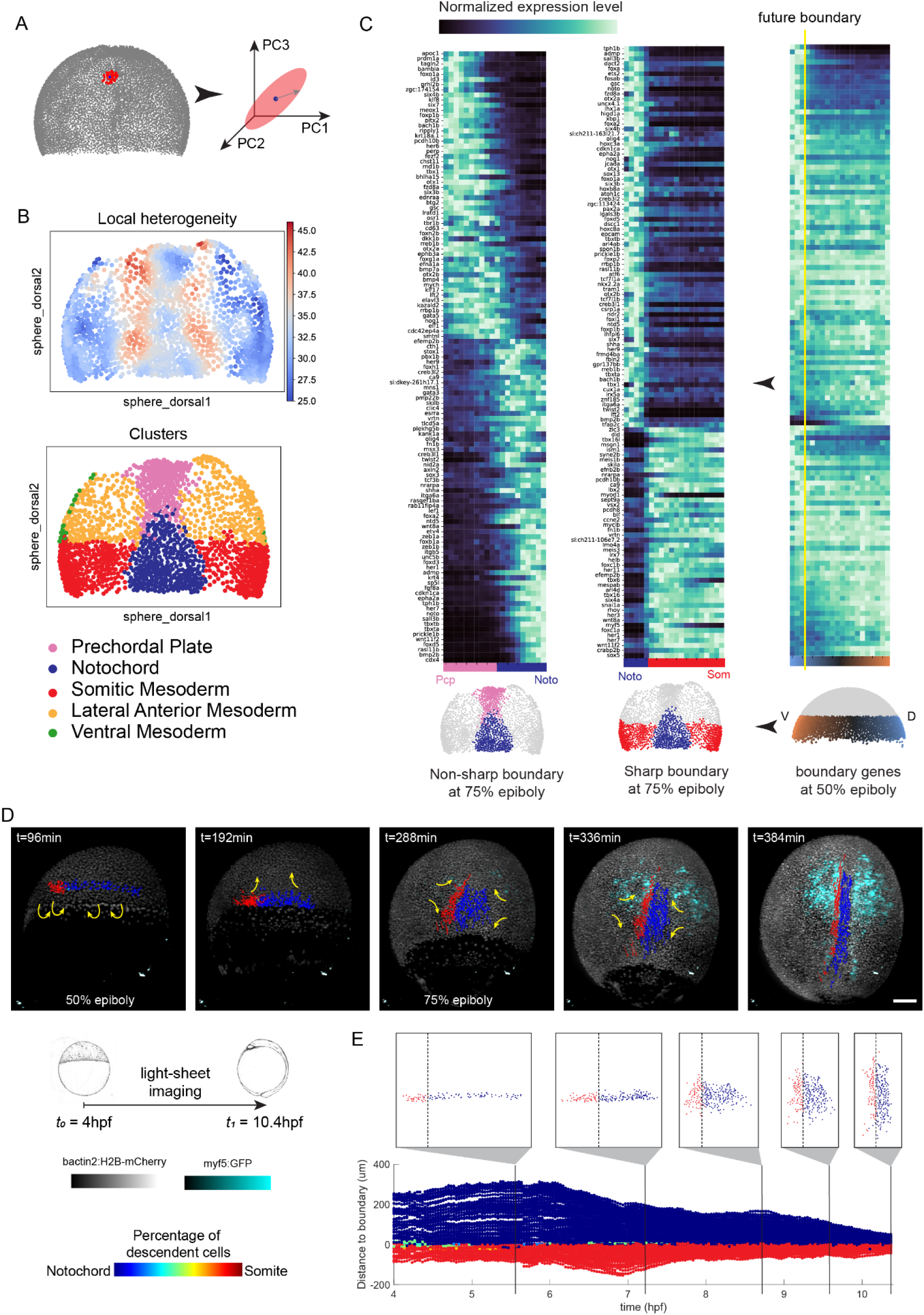
Sharp boundaries formed by staggered or aligned gene expression patterns defined by transcriptional regulation. **(A-B)** Quantification of local transcriptional heterogeneity: (A) Local transcriptional heterogeneity was defined for cells within each 50-μm neighborhood as the standard deviation in principal component analysis (PCA) space of their combined transcriptomes. (B) Heterogeneity map for the dorsal mesoderm shows a sharp boundary between the notochord and somite, contrasting with a non-sharp boundary between the notochord and the prechordal plate; **(C)** Left: Gene expression across sharp (notochord-somite) and non-sharp (prechordal plate-notochord) boundaries at 75% epiboly stage. Right: Gene expression along the dorsal-ventral direction within the notochord-somite progenitor cells at 50% epiboly stage; **(D)** Time-lapse imaging using light sheet microscopy shows boundary formation between notochord and somite. Nuclear mCherry marks all nuclei (gray) and myf5 highlights the muscle cells (cyan). The fates of cells marked in red for one side of the somitic mesoderm and blue for notochord are inferred from the last stages where myf5 positive cells are assigned as red and myf5 negative cells surrounded by myf5 expression are assigned as blue. The paths of cell movement (internalization and convergent extension) are shown in yellow arrows. Schematic illustration of the embryos are adapted from (*68*) **(E)** Quantification of distance to boundary over time of cells with notochord or somitic mesoderm fates. The graph shows minimal mixing between the progenitor cells of the notochord and somites.

To analyze the emergence of the sharp gene expression patterns observed at the notochord-somite boundary at mid-gastrulation, we analyzed their expression at the onset of gastrulation. We discovered that only a subset of genes have clearly delineated expression boundaries, while the rest are broadly expressed in the domain where notochord and somite precursors are located (Fig. 7C). Gene expression boundaries can emerge through multiple mechanisms, ranging from the sorting of cells of different types or lineages to the repression and activation of gene expression patterns without cell sorting (*50*). To determine the relationship between boundary formation, gene expression, and cell movement, we tracked notochord and somite progenitors from the blastula to the late gastrula stage using light-sheet microscopy (Fig. 7D). Cells were tracked using the ubiquitous nuclear-localized H2B-mCherry, and myf5:GFP was used to identify the cell fate at the somite-notochord border at the end of the time-lapse movie. We found that lineage restriction occurs early in development: at the onset of gastrulation (sphere stage), only 5 out of the 56 lineages consist of cells of mixed fate, and at mid-gastrulation stage (75% epiboly, 8hpf), each of the progenitors gives rise to either only notochord or only somite cells (Fig. S21A). Strikingly, there was very little cell mixing in the movement patterns of somite and notochord progenitors from blastula stage to mid-gastrulation (Fig. 7E) – fewer than 4% of cells crossed the boundary between the two progenitor types (Fig. S21B). The comparison of cell movement with gene expression indicates that genes co-expressed at the onset of gastrulation undergo a coordinated change in their expression to generate the sharp expression boundary observed at mid-gastrulation. This finding contrasts with the alternative hypothesis of the boundary arising from unmixing or cell sorting of a heterogeneous group of cells with different expression patterns. These results highlight the power of comparing multi-stage in situ spatial transcriptomics data with *in vivo* cell tracking data.

## Discussion

We introduce weMERFISH - an improved spatial transcriptomics method with subcellular resolution for profiling entire embryos in 3D - and establish a computational pipeline to integrate multiplexed imaging with scMultiome RNA and chromatin accessibility patterns into a common spatial atlas. We compiled six whole-embryo datasets, spanning stages from gastrulation to early organogenesis and mined these datasets for insights into the spatial regulation of the transcriptome across early embryogenesis. The study makes three major contributions: 1) a methodological development in both imaging and computational methods; 2) a comprehensive spatial transcriptomics resource; and 3) novel insights into the diversity, precision and emergence of embryonic patterns.

We overcame several of the limitations of existing spatial-omics methods and established an imaging platform to profile spatial gene expression for hundreds of genes in whole embryos at subcellular resolution. weMERFISH extends current approaches by enabling robust detection of transcripts in a multiplexed manner in thick 3D tissues. Compared to sequencing-based spatial transcriptomics (ST) methods (Visium, Stereo-seq (*51*), Slide-seq (*52*), Open-ST(*53*)), weMERFISH provides high subcellular resolution, high detection efficiency, and captures every single cell in the entire zebrafish embryo in 3D. Compared to cycleHCR (*54*), a recently pre-printed imaging-based spatial transcriptomics method that is 3D-compatible, weMERFISH is a highly multiplexed imaging method that can cover a large number of genes in fewer hybridization rounds. Compared to 3D MERFISH (*55*), which employed a similar confocal-imaging approach but utilized deep learning for *in silico* signal enhancement, weMERFISH uses branching signal amplification to physically amplify the signal, and by applying a flexible codebook constitution regime, offers the capacity to rapidly optimize the combinatorial imaging and validate it with single gene smFISH in the same sample. Instead of targeting a fixed gene panel, biologists can design a library of hundreds to thousands of the genes they are potentially interested in. They can decide later (even after sample preparation) to read out single genes using sequential imaging or to multiplex a large gene panel using a custom MERFISH codebook without the delay and cost of reordering the probes. weMERFISH is easy and cost-effective to implement, and broadly applicable to other systems such as *Drosophila* embryos or brains, whole zebrafish larvae, organoids, and thick tissue samples at the scale of hundreds of micrometers.

We constructed a resource of 6 embryos spanning from gastrulation to organogenesis with measured transcriptomes of 495 genes, imputed transcriptome of more than 20,000 genes and the accessibility patterns of more than 250,000 chromatin regions. The spatial information allowed the subdivision of previously discovered cell types and states and defined their precise spatial context. We make this dataset available and easy to explore with the aid of a beta-version web browser-based interface MERFISHEYES. The MERFISHEYES website contains the expression of all annotated genes and the spatial location of all accessible regions and will help users explore the expression and putative regulation of any gene of interest. Importantly, since gene expression is mapped in single embryos, the co-expression of genes can be analyzed at unprecedented accuracy. MERFISHEYES complements other websites such as ZFIN (*6*), gene expression atlas (*28*), DANIO-CODE (*56*) and Daniocell (*57*) and extends them to include high-resolution spatial information. It can be used to study cell-cell signaling, identify candidate genes in particular domains, or identify candidate enhancers with tissue-specific expression patterns.

We used the weMERFISH atlas to illustrate its potential in addressing two fundamental questions in developmental biology. First, what is the diversity and precision of gene expression patterns during embryogenesis? We found numerous novel gene expression patterns (Fig. S6), ranging from subcellular to global patterns. We found that transcripts have a preferential localization within cells and identify mRNAs associated with the cell periphery. At the tissue scale, we observe close to single-cell resolution positional information along the anterior-posterior axis and during border formation. At the global scale we discovered a large cohort of transcripts that are suppressed at the gastrula margin, a region that has been considered to be defined by the activation (not repression) of genes by mesendoderm inducers such as Nodal and FGF. It is unclear why factors involved in mitotic DNA replication initiation, chromatin organization and cell cycle are expressed at lower levels at the margin or how their repression might be regulated, but intriguingly, previous studies have suggested lower proliferation rates of cells located at the margin (*58*). Similarly, our analysis discovered a zone of cell-cycle inhibition in the emerging notochord domain. This region might serve as a “cell-cycle inhibition zone” that separates differentiating from proliferating cells, potentially allowing anterior proliferation waves to further differentiate the anterior notochord and the posterior proliferation wave to extend the tail (*59*). These observations lay the foundation for future functional studies. At the genome scale, the comprehensive mapping of gene expression domains defined genomic islands of co-expressed genes, revealed that co-expression in two or more tissues is frequent, and that co-expression is often observed in cell types that share functional or lineage properties. The analysis of accessible chromatin regions suggests that the diversity of expression patterns is generated by a combination of dedicated control elements, as in *Drosophila* (*60–62*), but also revealed more broadly accessible chromatin regions next to genes expressed in related cell types. Future mutagenesis and transgenesis studies will explore the roles of these putative regulatory regions in controlling gene expression.

Second, how do gene expression patterns emerge during embryogenesis? The combination of *in vivo* cell tracking and weMERFISH revealed the gene expression changes during somite-notochord border formation. While cells at the future somite-notochord border are spatially allocated already at the late blastula stage, most gene expression patterns found at the onset of gastrulation change dramatically to become restricted to either the somite or notochord territory at mid-gastrulation. Hence, sharp gene expression boundaries in this system do not emerge through cellular re-arrangements as observed in the spinal cord (*63*), but through changes in gene expression. How these gene expression patterns evolve or are regulated is unclear, but it is intriguing that we find staggered expression patterns spanning the developing notochord-prechordal plate border, and the dorsal and lateral posterior ectoderm border. It is possible that staggered patterns emerge as the notochord-somite border forms between the onset of gastrulation and mid-gastrulation. The staggered gene expression indicates that in a developing boundary, the position of a cell precisely determines its transcriptional profile, possibly gained from graded signaling pathways (*64–66*). In addition to cell tracking, we also used more indirect approaches to link spatial and temporal gene expression patterns. A limitation in our data is the lack of true temporal information on how gene expression changes within the embryo. To partially compensate for this limitation, we used computational tools to infer temporal information either by integrating the spatial data with existing molecular trajectories or through RNA-velocity, a nuclear/cytoplasmic RNA quantification method. The application of RNA-velocity generated vector fields that share the direction of morphogenetic movements and cellular differentiation. In the case of the internalizing margin, these vector fields might correspond to the spread of Nodal and FGF morphogen signaling, and in the case of cells converging to the midline, the vector fields might be caused by graded BMP signaling. The integration of temporal trajectories with spatial data illustrates the relationship of the gastrula margin with the tail bud and differentiating somites (Fig.4C), and of the axial mesoderm and differentiating notochord (Fig. S16). It is important to emphasize that these maps resemble specification maps that highlight the molecular state of cells, but do not replace fate maps and lineage tracing efforts. Future studies can build on these observations to determine the detailed connection between cell movements and gene expression dynamics.

In summary, the weMERFISH platform, consisting of the analysis pipeline, the data resource of spatial gene expression from gastrulation to organogenesis, and the beta version of the MERFISHEYES browser application extends and supports the genome-wide analysis of gene expression and regulation during embryogenesis. Our exploration of these datasets raises intriguing questions about the relationships between gene expression, border formation, cell maturation and morphogenesis, and lays the foundation for the future integration of multiple modalities into dynamic atlases of development.

## Supporting information

DataS1

DataS2

DataS3

DataS4

## Acknowledgements

We thank Long Cai, Simone Schindler, and Elsy Buitrago-Delgado for discussions on methodology development; Oliver Biehlmaier and the Biozentrum Imaging Core Facility for suggestions on instrumentation; Katarzyna Buczak for operating the liquid handler robot for linker probe reconstitution; Alba Aparicio Fernandez, Rita Gonzalez and Diana Medeiros Gomes for zebrafish husbandry. We thank Corinne Houart for critical reading of the manuscript.

This project has received funding from the Allen Discovery Center for Cell Lineage Tracing and an H2020 ERC Advanced grant (ERC-2018-ADG, grant agreement No 834788) to A.F.S. EMBO Long-Term Postdoctoral Fellowship (ALTF 709-2020) and an H2020 Marie Skłodowska-Curie grant (grant agreement No 101031809) to Y.W., Boehringer-Ingelheim-Fonds PhD Fellowships to J.E. and M.C.-T., and a research grant from NIH DP5-OD030878 to B.B.

## Author contributions

Y.W., B.B. and A.F.S. conceived and designed the study. Y.W. collected the data and performed the analysis on imaging and transcriptomics data. Y.W., J.E. and B.B. developed weMERFISH method and data processing pipeline with assistance from J.N.A. I.J. developed the MERFISHEYES browser website with assistance from E.L. and T-H.C. M. C.-R. performed the chromatin accessibility analysis with assistance from J.L. L.D. assisted in gene selection of the weMERFISH library. M.C.-T. piloted the spatial chromatin accessibility analysis. M.W. performed automated cell tracking of the light-sheet imaging data with supervision from G.Y. A.S. and S.E.M assisted the instrumentation of the weMERFISH imaging platform. Y.W., B.B. and A.F.S interpreted the results and wrote the manuscript with contribution from all authors. All authors read and approved the manuscript.

## Materials and Methods

### Gene Selection

We aimed to select ∼500 genes that are spatially and temporally varying from gastrula (50% epiboly) to early organogenesis (6-somite) stage. The initial candidate gene pool was selected from two sequencing datasets: a scRNA-seq datasets (*25*) and scMultiome dataset (*26*). For the scRNA-seq dataset, cell type identity and pseudotime have already been defined using URD (*25*). For the scMultiome dataset, cell type identity was defined by leiden clustering, and pseudotime was estimated by connecting each cluster along the most probable developmental trajectory using SlingShot (*69*). For both datasets earlier than bud stage, cell clusters in each segment of the pseudo-time trajectory were extracted and their enriched genes were identified and included in the candidate gene pool. Furthermore, along each trajectory, the top 100 variable genes correlating with pseudotime were also included in the candidate gene pool summing to n=1847 genes. Out of this pool we prioritized all annotated transcription factors (n=217) in the final gene list. Previously described marker genes were added to the final gene list, including marker genes from (*20*), markers for germlines, EVL, YSL, DFC and apoptotic cells (n=10), markers for each tissue type at 6-somite stage (n=39, DataS1) as well as cell cycle marker genes (n=11). Additional genes were selected from the candidate pool using an optimization algorithm that randomly selects groups of 1000-genes and checks how well the reduced gene set preserves the cluster identity along different stages in pseudotime (Fig. S2A). The top 163 variable genes out of the optimization results were added to the previous list (n=337) totaling 500 genes chosen for the weMERFISH imaging. Two metrics were used to quantify the ability of the selected genes to reconstruct developmental trajectories from the original scRNA-Seq data.

First, a k-NN graph was constructed with only the selected genes for a random set of N=1000 single-cells. For the k-neighbors of each selected cell, the neighbor cells’ deviation in assigned cell type identity (Fig. S2B) and deviation in pseudotime (Fig.S2C) was quantified. Both the 1000-gene list and the final 500-gene for weMERFISH can predict trajectory identity and pseudotime with comparable or even better performance than the full candidate pool. Second, using reduced gene sets we checked nearest neighbor cells’ cluster identity with the query cell cluster identity. Both the optimized 1000- and the final 500-gene libraries preserved the cell identity well (Fig. S2D-F). The UMAP structure of the original dataset is also preserved in the 1000- and 500-gene library (Fig. S2G-I).

### Primary probe design and preparation

Probe design is implemented in custom python script and available on GitHub: https://github.com/yinan-wan0/weMERFISH in the *1ProbeDesign* section. The primary probe set was constructed as previously described (*70*) and contained 5-60 40-mer target sequences complementary to the mRNA of each gene. Briefly, target sequence design included the following steps: (1) Building a 17-nucleotide hash-table with all the counts of each 17-mers as they appear in the unspliced transcriptome; (2) Scanning each 40-mer candidate sequence for each gene’s transcript and calculating off-targets based on the all 17-nt counts in the previous table; and (3) filtering 40-mer candidates based on predefined selection criteria (found in *1ProbeDesign/GenerateSequentialLibrary.ipynb*). We used the transcript sequences derived from the zebrafish reference genome sequences (GRCz11) downloaded from ncbi_refs eq. 5 genes were determined to be too short or too similar to another genes and 495 genes were selected for weMERFISH (Supplementary Data S2). Three 20nt readout sequences, unique for each gene, were concatenated to the 40-mer probes and two additional 20nt PCR primers were added on each end of the probe. These readout sequences and PCR primers were selected as previously (*70*) and screened against the Zebrafish transcriptome. The generation of the primary probe sets were prepared from oligonucleotide pools, as previously described (*14*). In brief, we used limited-cycle PCR to amplify the oligopools (Twist Biosciences) and cleaned up the PCR product using DNA oligo purification kits (Zymo Research, D4003). Then, we used these DNA sequences as the templates for in vitro transcription into RNA using T7 polymerase (NEB, E2040S). Subsequently, the RNA products were converted into single-stranded DNA with Maxima Reverse Transcriptase (Thermo Scientific, EP0751), and then the DNA was purified by alkaline hydrolysis (to remove the RNA templates) followed by DNA oligo purification kits (Zymo Research, D4006). The only difference is that the forward primer in the limit-cycle PCR as well as the reverse transcription steps, we use a 5’-acrydite modified primer, so that the primary probes can be covalently incorporated in acrylamide gels.

### Antibody conjugation

Goat anti-rabbit (Invitrogen, #31210) and donkey anti-rabbit (Invitrogen, A16019) were conjugated to DNA oligos containing linker sequences orthogonal to the RNA-targeting library as previously described in (*71*) with the modifications described in (*17*).

### Linker probe design and MERFISH codebook constitution

Linker sequences are gene-specific, which consist of a 20nt sequence reverse complimentary to the readout sequence of the genes’ primary probes, plus 2 repeats of amplification probe binding sites (20nt each). Linker probes were ordered from IDT in 384-well plates.

The MERFISH codebook was designed using a custom-written python script deposited at *1ProbeDesign/DesignMERFISHCodesZebrafish.ipynb*. Upon selecting the number N of hybridizations, for each color channel a set of binary words with 4 ones out of N was first randomly selected for each gene ensuring that the hamming distance between each pair of words is at least 4. This set of binary words associated to each gene dictates in which hybridization each gene should appear and is called the MERFISH codebook. An optimization algorithm based on Metropolis Hastings (*72*) was used to swap the binary words in the codebook associated with each gene in order to minimize the average crowding in each hybridization of the transcripts expressed in each cell type as inferred from scRNA-seq.

The optimized MERFISH codebook was constituted using a pipetting robot (Tecan Fluent) that pipetted out from 384-well plates the linker for each gene and mixed it according to the MERFISH codebook into 1.5 mL DNA Lo-bind Eppendorf tubes corresponding to each round of hybridization.

### Amplification and readout probe design

Level-1 amplification probes consist of a 20-nt arm that complements the overhang of the linkers, and an additional overhang of 4x binding sites (20-nt each) for the level-2 amplification probe. Similarly, level-2 amplification probe consists of 20-nt that binds to level-1 amplification overhangs, and 4x binding sites (20-nt each) for the fluorescent readout probes. Amplification probes were ordered from IDT.

Readout probes are 5’- and 3’-dye-modified oligos of 20-nt length. The dye modification is Cy3 for genes imaged with 561nm laser, and Cy5 for genes imaged with 638nm laser, and Alexa Flour 488 for genes imaged with 488nm laser. Readout probes were ordered from Eurofins Scientific with HPLC purification.

The sequences of the amplification and readout probes can be found in Supplementary Data S3.

### Zebrafish

Embryos from wild-type (TL/AB) crosses were collected 20 minutes after fertilization. They were raised in E3 medium at 28.5°C until the designated stages based on hours post fertilization (hpf)(*68*). Once reaching the designated stage, the embryos were fixed within the chorions in 4% PFA at 4°C overnight, dechorinated manually using forceps after washing with PBS, and subsequently dehydrated with a gradient concentration of methanol in PBS solution (100%PBS – PBS:methoanol 1:1 – 100% methanol) and stored at -20°C for up to 1 month. Zebrafish sex cannot be determined until ∼3 weeks post fertilization, and thus the sex of experimental animals was unknown.

### Sample preparation

Frozen dehydrated zebrafish embryos were rehydrated with reverse methanol gradient (PBS:methoanol 1:1 – 100% PBS). The embryos were removed from the yolk. We first performed antibody staining on the rehydrated embryos: samples were incubated in blocking solution (1X PBS, 0.25% Triton X-100, 10mg/ml BSA, 0.5 mg/ml salmon sperm DNA, 1:2000 RNAse Inhibitor) for 2 hours at room temperature, then incubated with primary antibodies diluted in blocking solution at 4°C overnight. Here we used mouse anti-e-cadherin (BD Biosciences, #610181, 1:500 dilution) and rabbit anti-beta-catenin (Sigma-Aldrich C2206, 1:250 dilution). The samples were washed three times with PBS-0.1% Triton X-100 (PBST), and then incubated in conjugated secondary antibodies in blocking solution (concentration 10 µg/ml) for 2 hours at room temperature. The samples were washed three times with PBST and postfixed with 4% PFA for 15min at room temperature, then washed three times with 2xSSC + 0.1%TritonX-100 (2xSSCT).

For primary probe hybridization, embryos were incubated in pre-hybridization solution (40% formamide in 2xSSC-T, 1:500 RNase inhibitor) for 30 minutes at room temperature then incubated in hybridization solution (50% dextran sulphate, 50% formamide in 2xSSCT, 1:100 RNase inhibitor) at 47°C from overnight to 2 days. Embryos were washed in 40% formamide in 2xSSCT for 1 hours, then rinsed twice with 2xSSCT for embedding.

Coverslip functionalization and gel polymerization was performed as described in (*16*). Embryos were pre-treated with hydrogel solution without polymerizers (4% A/BA stock, 0.3M NaCl, 0.06M Tris-HCl, 1:5000 RNase inhibitor) overnight at room temperature. They were subsequently mounted to the coverslip with the EVL cells facing the coverslip, with 75µL hydrogel solution plus 0.05% (w/v) APS and 0.05% (v/v) TEMED. A cut is made into the embryos in order to unravel them onto the coverslip. The hydrogel was left in hydrogen chamber for polymerization for at least 2 hours at room temperature, sandwiched between a GelSlick-treated large glass slide. The coverslip was gently peeled off the coverslip and set in a plastic petridish. The slide was cleared with 1.25% (v/v) SDS and 0.2 mg/mL proteinase K solution in 2xSSC at 47°C for 3 hours, postfixed with 4% PFA for 15min at room temperature, and proceed to imaging.

### MERFISH imaging with signal amplification

Imaging was performed on a Nikon Ti2 with CSU-W1 spinning disk confocal microscope (50um pinhole single disk), coupled to a home-built fluidics system similar to one previously described for MERFISH imaging (*73, 74*). We coordinated imaging and the fluidics system control using custom macros and the Nikon Elements software. Imaging was achieved using the Nikon CFI Plan Apochromat VC WI 60x/NA 1.2 objective. A 405-nm laser was used to excite the nuclear stain DAPI, providing signal for registration. Two laser colors were used to image the genes combinatorially in the codebook or sequentially for the genes outside codebook or for ground truth: 561nm laser for the Cy3 genes and 638nm for the Cy5 genes, and the detection was through a quad-band emission filter (Semrock FF01-440/521/607/700-25). Fluorescence was imaged with a CMOS camera (Hamamatsu Fusion). Each FOV was imaged at a single z-plane with a 300-500ms-exposure time for the Cy3/Cy5 channel, and 100ms for the DAPI channel. Samples were hybridized with 1) linker, 2) level-1 amplification, 3) level-2 amplification and 4) fluorescent readout probes and imaged following protocols similar to those previously described (*14*). Probe hybridization buffers were composed of 2xSSC-T, 35% (v/v) formamide and 25nM (for combinatorial linkers, concentration per probe), 100nM (for sequential linker probes and amplification probes) or 200nM (for fluorescent readout probes) of the appropriate probes. 1mL hybridization probe buffer was used per imaging round, and excessive probes were washed out using 5mL wash buffer composed of 2xSSC-T with 30% (v/v) formamide between hybridization rounds. After hybridization and washing, 2mL imaging buffer was flown into the imaging chamber containing 2× SSC, 50 mM Tris⋅HCl pH 8, 10% (m/v) glucose, 2 mM Trolox (Sigma-Aldrich, 238813), 0.5 mg/mL glucose oxidase (Sigma-Aldrich, G2133), 40 μg/mL catalase (Sigma-Aldrich, C3155), and 4ug/mL DAPI (Sigma-Aldrich, D9542) solution, after which the flow was halted and 43∼47 FOVs were imaged, each consisting of ∼300 z-slices at 0.3 um z-step size. The z-location of the sample was kept stable using Nikon’s Perfect Focusing System. After each round of imaging, the linker, amplification and readout probes were stripped using 5mL of 80-100% (v/v) formamide, only leaving the primary probes in the sample.

After the MERFISH rounds of imaging, sequential genes were imaged followed by a blank bit and a series of genes in the codebook readout sequentially as a ground truth, and the sequential genes not in the MERFISH codebook. Finally, the membrane-conjugated antibodies were imaged at 405 or 561nm, as sometimes the antibody signal can get very bright and hard to strip.

### Image processing

Processing of the images was conducted using custom-written python code available on GitHub: https://github.com/yinan-wan0/weMERFISH, folder “2ImageProcessing”. More specifically, the code can be broken down into 4 steps.

#### 1. Dot detection

Raw images of the genes in Cy3/Cy5 channel were first subject to flat-field correction. Then the images were deconvolved using the PSF (point spread function) subtracted from previous images. The images were then normalized and detected for Gaussian peaks using the parameter of sigmaXY=1.85 pixels, sigmaZ=2.5 pixels, and a minimum intensity threshold of 80. The code also output each dot’s intensity in raw and deconvoluted space, correlation with Gaussian in raw and deconvoluted space for further filtering.

#### 2. Image registration

Image registration consists of two parts: registration across hybridization rounds, and registration across FOVs.

For registration across hybridization rounds, the DAPI channel was deconvolved and detected for features of local maxima and minima points using the same detection algorithm as dot detection. Images were aligned across hybridization rounds by maximizing phase cross-correlation on the binarized maxima/minima feature point images. The feature extraction enhanced the texture inside the nuclei and provided a very robust way of translation estimation.

Registration between FOVs was only performed for the hybridization round of membrane antibody. An initial overlap (10%) was first extracted from the imaging settings, and image registration was achieved by maximizing the phase cross-correlation. Offsets between neighboring FOVs in the horizontal and vertical directions were calculated, and a global offset was further computed from the pairwise offsets.

#### 3. Cell segmentation

Cell segmentation consists of two parts using Cellpose (*27*): 3D segmentation of the deep cells, and 2D segmentation of the surface/EVL cells, as the latter have very flat morphology and are very hard to fit into the prediction of Cellpose’s isotropic 3D-segmentation model. First, cells were segmented using the DAPI and membrane channels using the Cellpose ‘cyto2’ model with a diameter of 33. Then, the cell nuclei were segmented using only the DAPI channel using the Cellpose model ‘nuclei’ with a diameter of 20. The two segmentation results were combined so that small segmentations that do not contain a nuclei were removed.

For segmenting the EVL cells, the DAPI and membrane channels were first subject to Cellpose’s 2D segmentation model ‘cyto2’ using a diameter of 50 pixels. Then, the dots corresponding to hybridization round for gene krt4 was extracted, and 2D masks with more than a threshold number of molecules of krt4 were defined as EVL. 2D segmentations on consecutive z-planes with intersect-over-union value of 0.5 were stitched in 3D. The EVL mask was expanded by 1 pixel in all directions (to address sometimes the discontinuous 2D segmentation) and combined with the rest of the 3D segmentation. This ended up giving ∼9,300 cells at 50% epiboly stage, and ∼18,200 cells at 75% epiboly stage.

For 6-somite stage data, oversegmentation was a problem for FOVs where the majority of space was empty, which led to an overestimation of number of cells compared to the reported number (*75*). To address the problem, the stitched DAPI image was again fed to Cellpose for a global segmentation. Segmented regions that contain the same nucleus were merged into a new cell, and segmented regions that do not contain any nuclei were removed. This ended up giving ∼24,200 cells at 6-somite stage.

Segmentation results for neighboring FOVs was consolidated in a way that cells closer to the center of the corresponding FOV were kept, and the cells further away from the center of the FOVs were discarded. A new unique global ID was given to each cell, and the images of the nuclei and membrane channels were stitched for global visualization.

#### 4. MERFISH decoding

The spot-based decoding was implemented as previously based on colocalizing fitted spots across all hybridizations within a distance threshold and then matching the brightest spots to the corresponding binary words in the MERFISH codebook (*15, 70*). One modification were implemented compared to the previous decoding: the decoding was done in 3-passes to initially use a generous co-localization threshold (6 pixels) to correct for distortion in z and local distortion in x,y and z, and finally decoded with a conservative threshold (2.5 pixels) to match the designated codebook. Molecules were filtered to remove potential false positives using 1) the correlation of each spot with a Gaussian point spread function and 2) the distance between the normalized brightness of the colocalizng spots of each decoded mRNA to the closest barcode. The filtering thresholds were selected such that the detection of blank barcodes (combination of spots that did not have a corresponding gene in the MERFISH codebook) was less than 7%. RNA molecules identified were assigned to cells by applying the segmentation masks to the positions of the molecules. The scripts for MERFISH decoding are deposited at *2ImageProcessing/MERFISH_PIPELINE.ipynb*

### Cell clustering analysis of MERFISH

Combining cell segmentation with the MERFISH single-molecule detection, we constructed cell-by-gene matrices. The matrices were used in combination with the Scanpy version 1.10 package to further analyze the MERFISH data. Cells with too few total transcripts were likely segmentation of autofluorescence and were therefore discarded. Count normalization, Principle Component Analysis (PCA), neighborhood graph construction and Uniform Manifold Approximation and Projection (UMAP) were performed with Scanpy’s default parameters. We performed Leiden clustering using multiple resolution parameters. The top 20 differential genes identified by the rank_gene_groups function and their physical locations were used to annotate each cluster. For 50% epiboly stage, annotation is done on one embryo and then transferred to another embryo using scanpy’s ingest function, for 75% epiboly stage embryos, joint annotation is done using batch correction, for 6-somite stage’s two embryos that were acquired with exactly the same condition, no batch correction was necessary. The scripts for cluster annotation can be found in the Github folder *3CellTypeAnnotation*

### Reconstruction of flattened data to 3D anatomy

For 50% and 75% epiboly stage data, the embryos that were unraveled from the spherical yolk were reassembled on a spherical shell using the following procedure:

A) The size of the 3D spherical shell onto which the embryos were reassembled was determined based the size of the margin and the area of each embryo. An optimization algorithm was implemented with the pytorch package to map points uniformly covering from the unraveled 2D view onto points on the 3D spherical shell. The algorithm satisfied the following constraints:

1) The margin of the embryo and the dorsal axis (marked in red in Fig.S4 A-B) were enforced to map from the 2D unraveled view onto the 3D view in fixed positions.
2) The two edges of the cut on the embryo which allowed for its unravelling (marked in green and blue in Fig.S4A-B) were stitched together.
3) The local distances between points in 2D were matched to the distances on the spherical shell in 3D. This mapping from 2D onto the 3D spherical shell was applied all the cells in the unraveled view for the reconstruction.

For 6-somite stage, to restore the flattened anatomy in the z direction, we acquired the anatomy of 3 intact zebrafish embryos of genotype Tg(bactin2-HRASGFP)Tg(bactin2-H2BmCherry) with ubiquitous membrane and nuclear markers at 6-somite stage with Zeiss Lightsheet 7 microscopy from two opposing side views. The data was reconstructed using Fiji BitStitcher plugin (*76*). The stitched image volume was visualized using the Imaris software (Bitplane inc) and the midline plane was fitted with oblique slicer (Fig. S4C). The body axis as well as the thickness of the sample at ∼30 reference locations were measured with the python napari plugin, and a curve of thickness as a function of the anterior-posterior (AP) location was fitted from 3 measured embryos (Fig. S4D) and used to rescale the flattened embryos.

### Integration of single-cell multiome and weMERFISH

To integrate the scMultiome and weMERFISH datasets, we utilized the software Tangram (*36*) that used non-convex optimization and a deep learning framework to learn a spatial alignment for the RNA sequencing data and our MERFISH measured genes. The mapping probability matrix from single-cell to spatial data was estimated by Tangram’s map_cells_to_space function ran in cells mode with a uniform density prior. Then the genes were imputed using Tangram’s project_genes function that multiplies the probability matrix with the single-cell transcriptome.

As for the single-cell ATAC-seq data, the Term Frequency-Inverse Document Frequency (TF-IDF) score for each detected peak was used as the input, and the imputed chromatin accessibility was similarly computed using tangram’s project_genes function. The scripts for gene imputation can be found in the GitHub folder *4MultiomeMapping*

### Modified RNA-velocity analyses

RNA velocity was calculated using the software scVelo (*46*), with nuclear and cytosolic count stored in to ‘spliced’ and ‘unspliced’ layers. After filtering and normalizing, velocity graph was generated using the deterministic model, and the velocities were projected into a lower-dimensional embedding both on UMAP and 2D flattened view of the embryo, with germlayers offset relative to each other. For 50% and 75% epiboly stage data, both spherical views and flattened views were used to embed the velocity stream.

### 6-somite stage Gene enrichment analysis

For the 6-somite stage gene enrichment analysis, only the 495 measured genes and 3377 highly variable genes in the original single-cell multi-omics dataset were considered. Genes enriched in different tissues were identified using Mann-Whitney test between the tissue of interest and the rest of the cells, with p-value threshold set to 1e-50 and log2 fold change threshold set to 1. Epidermis, YSL, PGC and EVL related tissues were removed from the analysis as they do not exist in the scMultiome data or they contain contaminated signals from cells nearby. To plot enriched genes in the brain, the tissues were divided anterior-posteriorly into 100 bins, and average gene expression in the corresponding bins were plotted as a heatmap.

### Dual-tissue accessibility analysis

We identified genes expressed within two tissues domains. Accessible chromatin peak regions that were within 100kb of the dual-tissue expressed genes were subject to further analysis. Peaks enriched in different tissues were identified using Mann-Whitney test, with p-value threshold set to 1e-50 and log2 fold change threshold set to 0.5.

We investigated the genomic locations of genes uniquely expressed in specific cell types across the 6-somite stage embryo. Gene density was assessed by examining the presence of other genes at various distances from the transcription start site (TSS) up to 500,000 base pairs. Genes enriched within the same tissue were classified as cell-type enriched, enriched in other cell types, or uniformly distributed across all cell types. To quantify this enrichment, we calculated the gain as the normalized density of enriched genes at a given genomic distance, divided by the normalized density observed between 300,000 and 500,000 base pairs from the TSS, which served as a control region. We then repeated the same procedure with ATAC-seq peaks, focusing on peaks located near genes expressed in a single tissue. We assessed whether these ATAC peaks were also enriched in the corresponding cell type, in other cell types, or were uniformly accessible across all cell types.

### Subcellular localization analysis

To measure the distance between the transcripts to cell membrane, we performed distance transform on the segmentation masks. Statistics were performed for the genes enriched in each cell type selected (e.g. EVL marker genes were excluded from distance analysis for deep cells). Data was plotted with scale bars as mean ± SEM across all transcripts of the specific cell type in an embryo.

### Boundary sharpness quantification

The sharpness between domains of gene expression corresponds to local heterogeneity. To quantify the local heterogeneity of gene expression, we focused on cells in the deep tissue and removed extraembryonic tissues for the analysis. We took cells from a local neighborhood (distance smaller than a threshold), and measured the spread of the neighborhood cell’s in the PCA space by averaging the L2 distance between each of the neighbor cell and the center of the neighborhood in the PCA space. The analysis was first ran on measured data, then repeated for imputed data. Multiple neighborhood sizes (20, 30, 40, 50 um) and number of PCs (50, 100, 200) were tested to ensure the robustness of the analysis result.

### Light-sheet imaging of boundary formation

Transgenic zebrafish embryos with fluorescent nuclei marker Tg(bactin2:H2BmCherry) (*77*) and Tg(myf5:GFP) (*78*), inside their chorions, were embedded in 1% low melting point agarose prepared in E3 medium, enclosed by glass capillary before extruded into the imaging chamber. Images were acquired with Zeiss LightSheet 7 Microscopy, with 20x/N.A. 1.0 detection objective (additional optical zoom factor 0.55x) and dual-side 10x/N.A. 0.3 illumination objectives. Fluorescence of GFP was activated by 488nm laser and detected with BP505-545 filter, and fluorescence of mCherry was activated by 561nm laser and detected with LP585 filter. Time-lapse imaging was performed at 2-minute interval from 4 to 12 hours post fertilization. Within each time interval, four 3D volumes were acquired with 90-degree rotation in between to achieve full-embryo multiview coverage. The z-stack was set to have the voxel size of 0.43 um x 0.43 um x 2.5 um, so that each cell nuclei is sectioned by at least 3 planes.

Automatic cell tracking was performed using the muSSP algorithm (*79*) in a custom-designed software written in MATLAB. The notochord and adaxial cells were identified at the end time point and back curated using the Fiji plugin mastodon (https://imagej.net/plugins/mastodon). To visualize the boundary, a Support-Vector-Machine (SVM) decision plane was fitted to every time point that best distinguishes the notochord- and adaxial-fated cells, and the distance to the plane was measured.

---

**Fig. S1.**
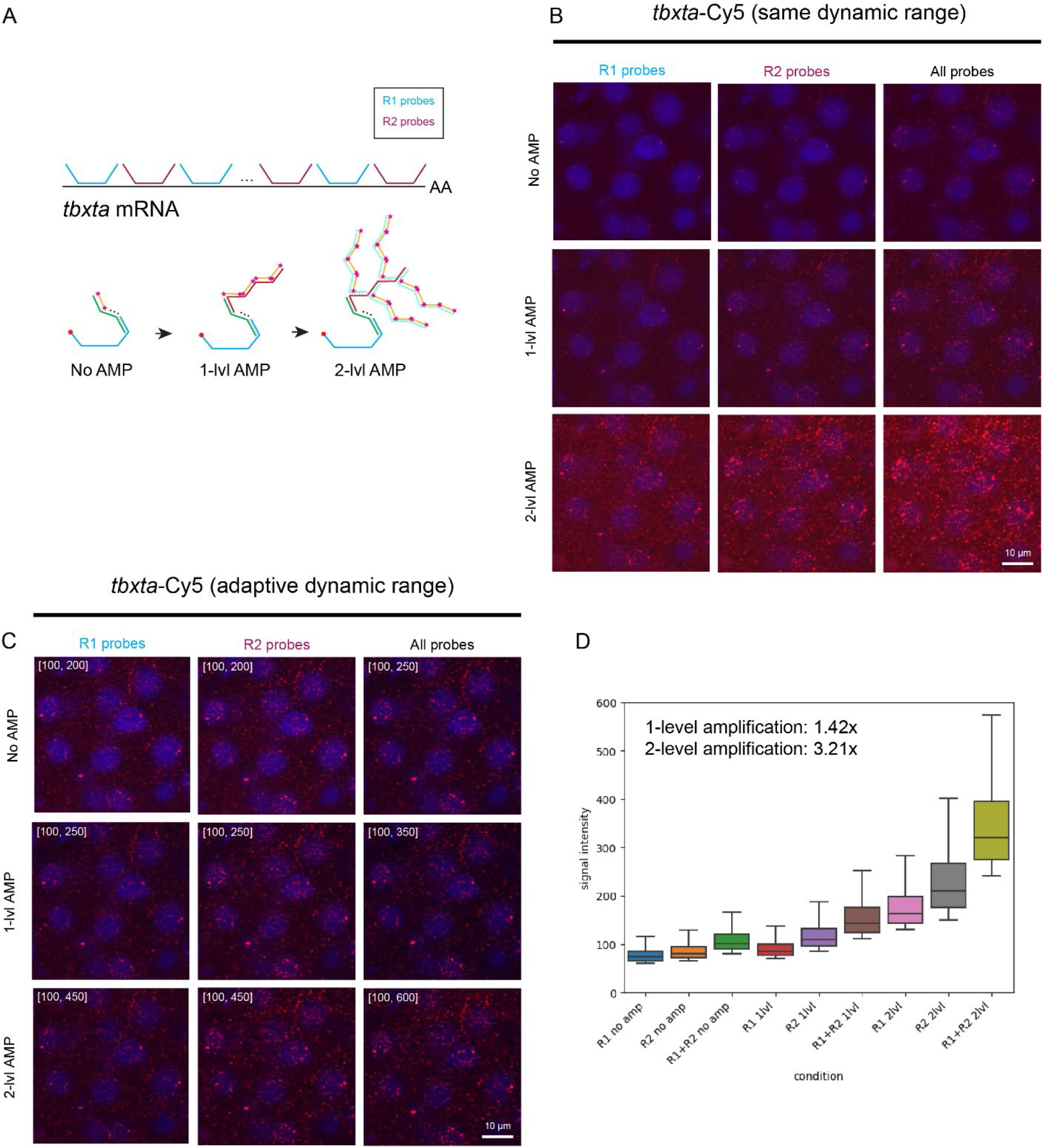
Branching amplification of the single-molecule signal. **(A)** Schematic of amplification process: The gene tbxta was selected for testing the amplification strategy and two sets of interleaving encoding probes were designed to target the transcript and have two readout probes R1 and R2. The amplification process starts with hybridization of the encoding probes, followed by 1-level amplification against R1 or R2 probes, then a 2-level amplification where additional probes increased signal branching and finally the readout probes with fluorescent dyes. **(B-C)** Amplification of tbxta-Cy5 signal: (A) Imaging of tbxta-Cy5 signal without amplification (No AMP), with 1-level amplification (1-lvl AMP), and with 2-level amplification (2-lvl AMP) under the same dynamic range. R1 probes (blue), R2 probes (pink), and combined probes are shown. (B) Images in (A) with adaptive dynamic range, adjusting the display intensity for each amplification level. Signal intensities are annotated for R1 and R2 probes, and the combined signal is shown in each row. **(D)** Quantification of signal intensity: Box plot showing signal intensity for different conditions (No AMP, 1-lvl AMP, 2-lvl AMP) for R1 and R2 probes. The average signal amplification is quantified, showing a 1.42x brightness increase with 1-level and a 3.21x increase with 2-level amplification.

**Fig. S2.**
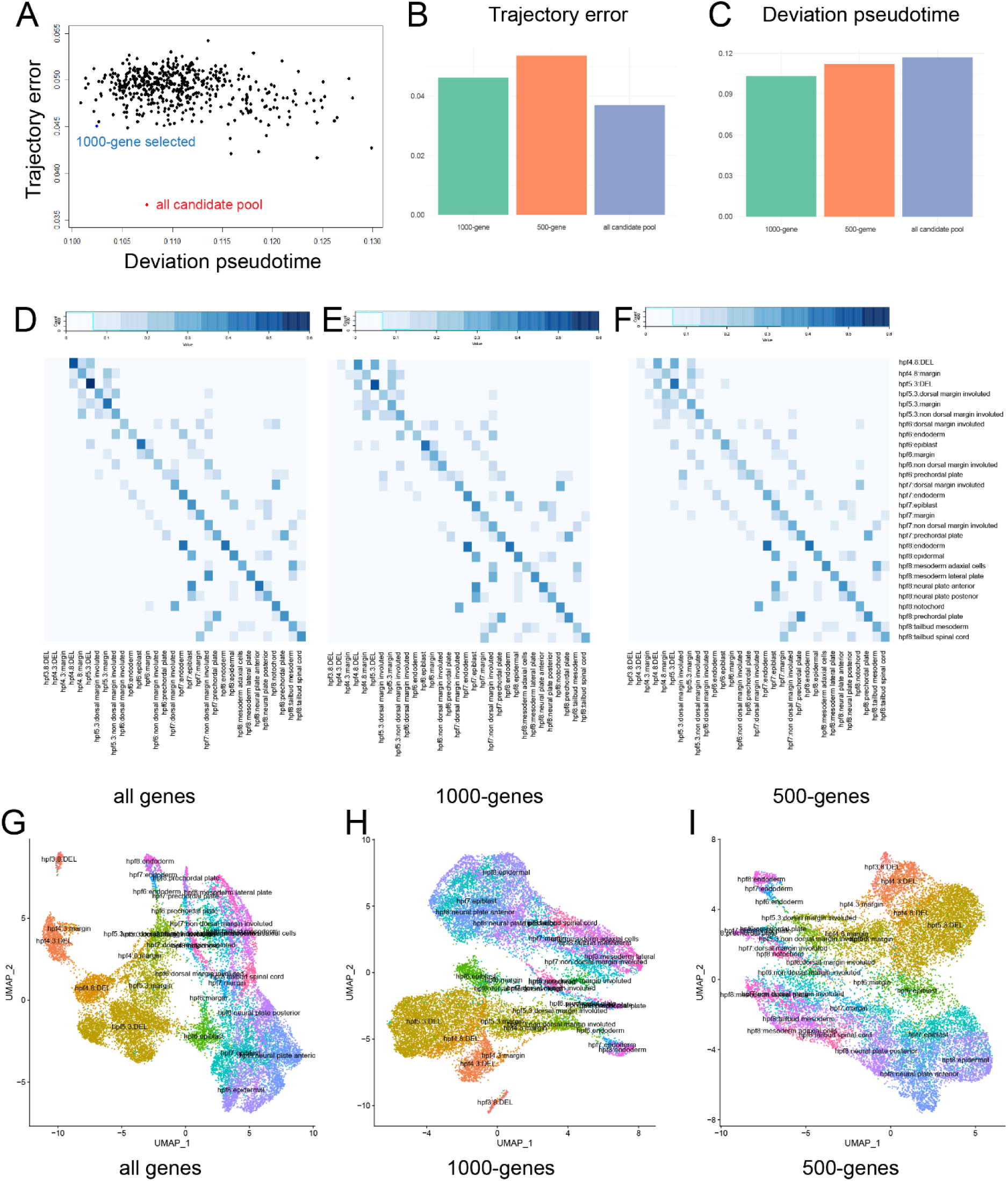
Selection of variable genes for weMERFISH. **(A)** Selection of subsets of 1,000 genes from the initial candidate pool using an optimization algorithm, evaluated for cell type error (called trajectory error) and deviation in pseudotime. The scatter plot correlates trajectory error and pseudotime deviation for selected genes (blue) compared to the entire candidate pool (red).; **(B-C)** Comparison of trajectory error and pseudotime deviation across an optimized 1,000-gene set, the 500-gene MERFISH set, and the full candidate pool. The bar plots indicate that both gene libraries maintain lower trajectory errors and pseudotime deviations. **(D-F)** Heatmaps showing the accuracy of predicting cell identity based on the 30 nearest neighbors using all genes, the 1,000-gene library, and the 500-gene library. The libraries effectively preserve cell type identification accuracy. **(G-I)** UMAP visualizations reconstructed using all genes, the 1,000-gene library, and the 500-gene library. The overall topology and cell type identities are well preserved across different gene libraries.

**Fig. S3.**
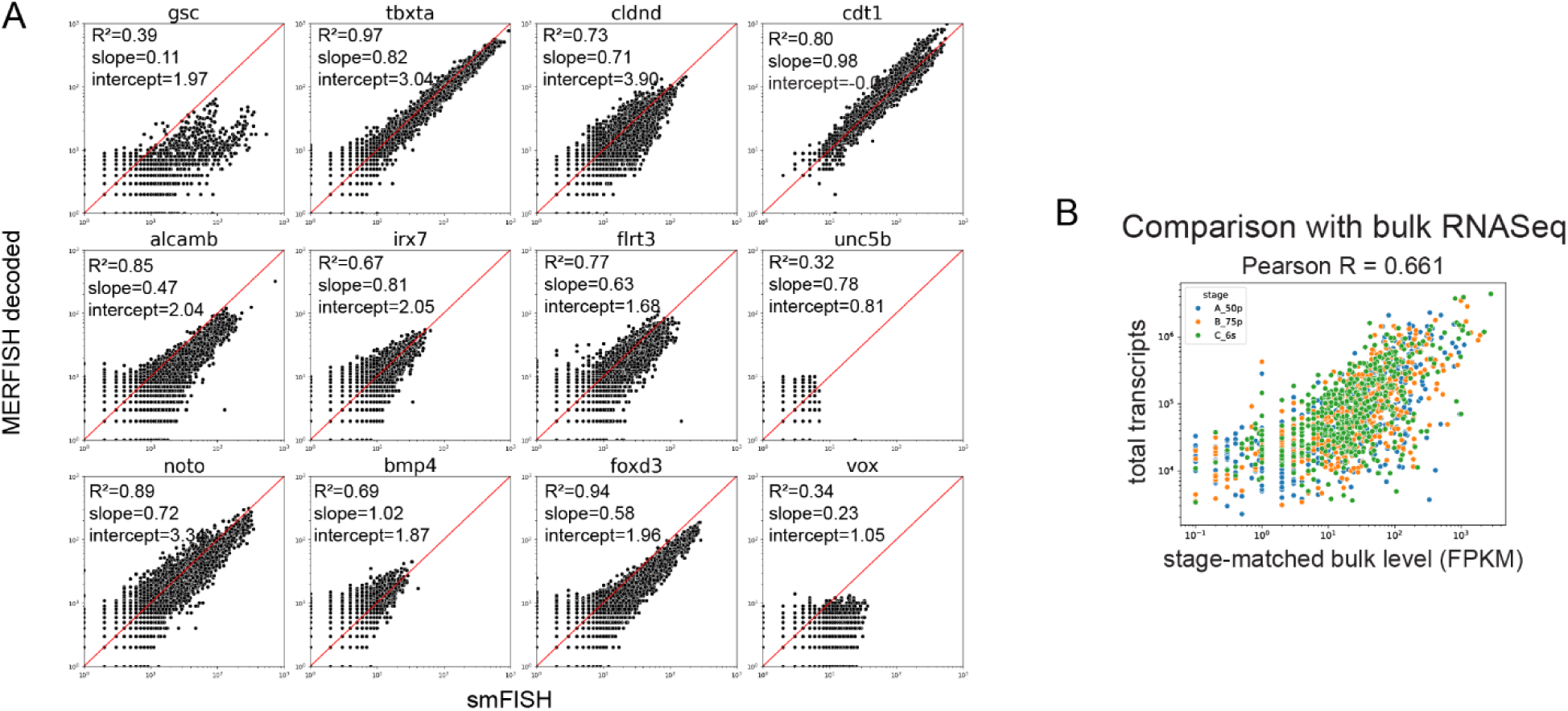
Quality control of weMERFISH measurements. **(A)** Comparison of weMERFISH quantification with smFISH molecule counts per cell at the 50% epiboly stage. Scatter plots for various genes (e.g., *gsc*, *tbxta*, *cdt1*) show the correlation between MERFISH and smFISH data, with R² values, slopes, and intercepts indicating the degree of agreement between the modalities **(B)** Correlation between total transcripts measured by weMERFISH and stage-matched bulk RNA-Seq across three developmental stages (Pearson correlation coefficient R = 0.661).

**Fig. S4.**
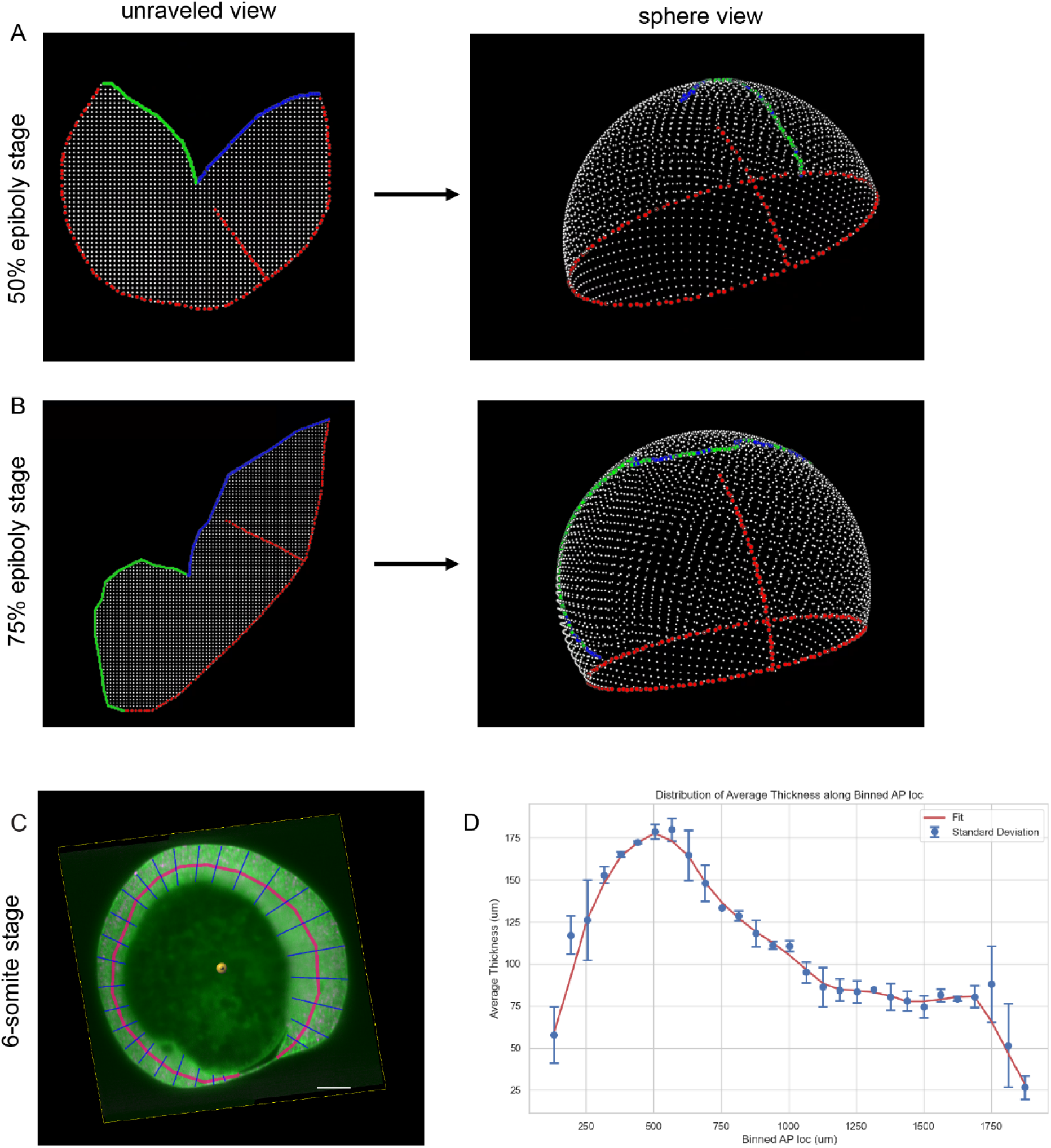
Reconstruction of the unraveled view to original anatomy view. **(A-B)** At 50% epiboly (A) and 75% epiboly (B), the unraveled 2D views of embryos are reconstructed into 3D spheres. The embryo’s margin and dorsal axis (red) are mapped to fixed positions in the 3D view, while the edges of the cut used for unravelling (green and blue) are stitched together. Local 2D distances between points are matched to the corresponding distances on the 3D spherical shell. (C) Light-sheet imaging of the midline plane of a 6-somite stage embryo is used to measure the body axis thickness. The image shows the measurement grid superimposed on the embryo. (D) Summary statistics of the body axis thickness across three 6-somite stage embryos. The thickness distribution is plotted along the anterior-posterior axis, with error bars representing standard deviation.

**Fig. S5.**
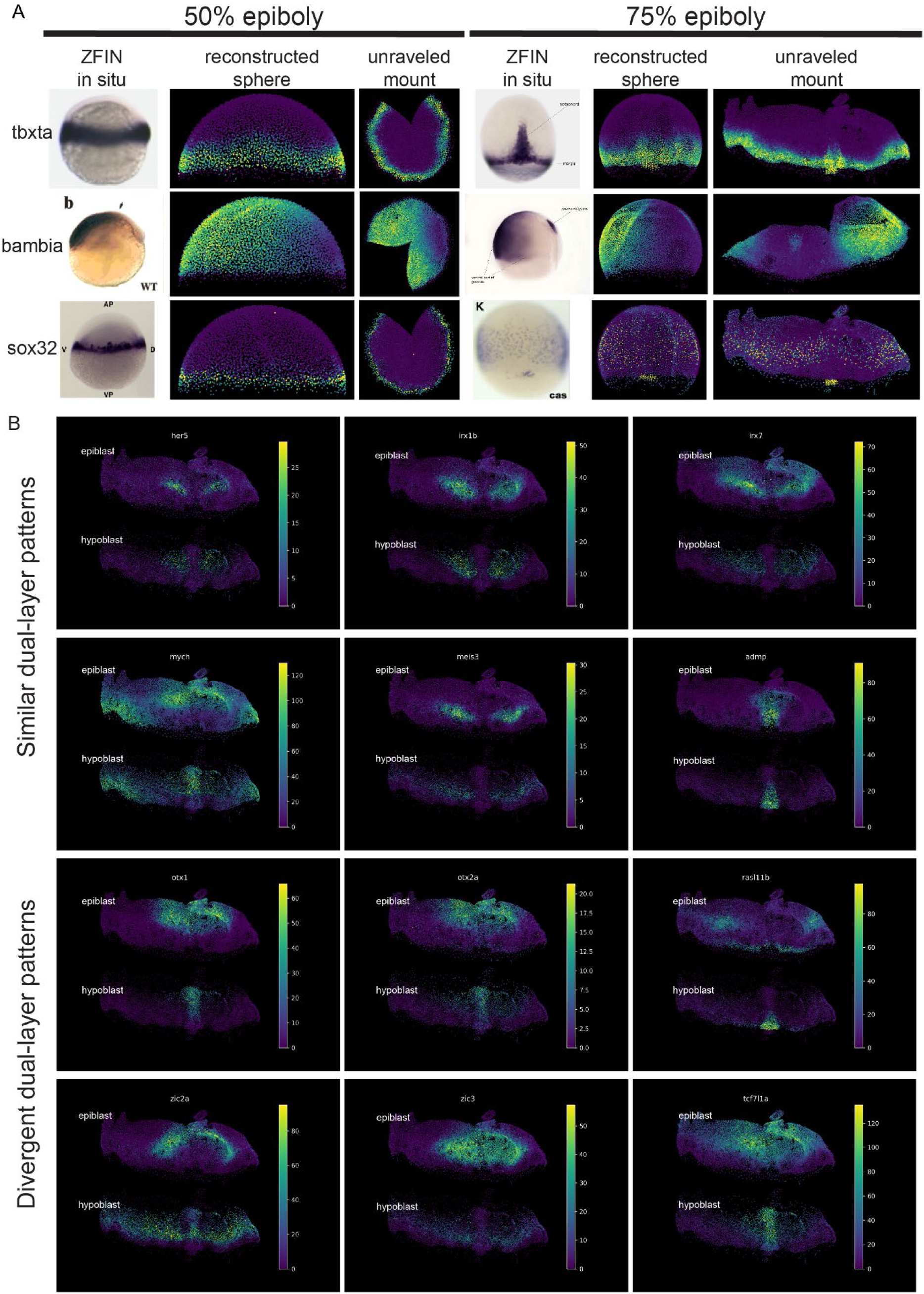
Expression patterns by weMERFISH are validated by and refine prior in situ hybridizationd work. **(A)** Comparison of weMERFISH data with published in-situ data: The left column shows *in situ* hybridization images from the Zebrafish Information Network (ZFIN) database for three genes (*tbxta*, *bambia*, *sox32*) at the 50% and 75% epiboly stages. The middle and right columns display the corresponding weMERFISH data in both reconstructed 3D spherical views and unraveled mounts, demonstrating consistency between the weMERFISH and *in situ* data. **(B)** Genes with dual-layer (epiblast and hypoblast) expression patterns at 75% epiboly stage: Visualization of genes exhibiting dual-layer expression in both the epiblast and hypoblast. Examples include: Similar dual-layer patterns: Genes like *her5*, *irx1b*, and *mych* show similar expression in both layers. Divergent dual-layer patterns: Genes like *otx1*, *etv2a*, and *tcf7l1a* exhibit differing expression levels between the epiblast and hypoblast. This figure highlights the validation of weMERFISH data against established *in situ* results and reveals complex, layer-specific gene expression patterns during zebrafish embryogenesis.

**Fig. S6.**
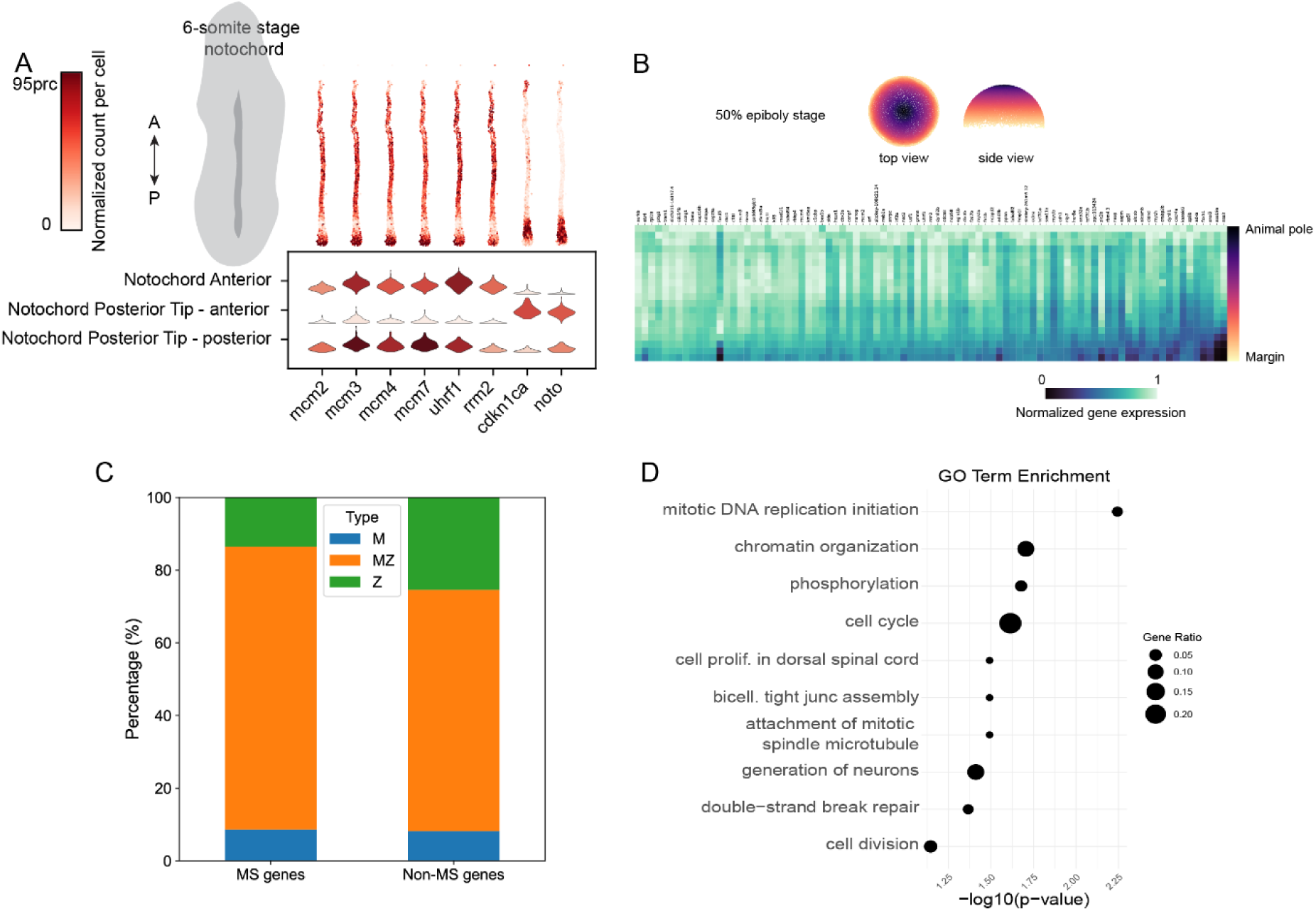
Novel gene expression patterns at the tissue and organismal levels. **(A)** Notochord cells at the 6-somite stage show varying expression levels of cell cycle genes across anterior (A) to posterior (P) subclusters. Violin plots depict normalized counts per cell for key genes such as *mcm2*, *mcm3*, *uhrf1*, and *noto*. (A) to posterior (P) subclusters. The violin plots show the normalized count per cell for key genes. **(B)** At the 50% epiboly stage, 83 genes are suppressed at the embryonic margin. The heatmap shows normalized gene expression, with suppressed genes localized toward the margin (colored in yellow) as opposed to higher expression at the animal pole (colored in purple). **(C)** Bar plot illustrating that marginally suppressed genes are primarily maternal-zygotic (MZ, orange) rather than maternally deposited (M, blue) or zygotic (Z, green). **(D)** Bubble plot of GO term enrichment for the marginally suppressed genes, highlighting processes such as mitotic DNA replication initiation, chromatin organization, cell cycle, and cell division, with gene ratio and statistical significance (p-value) indicated by bubble size and position.

**Fig. S7.**
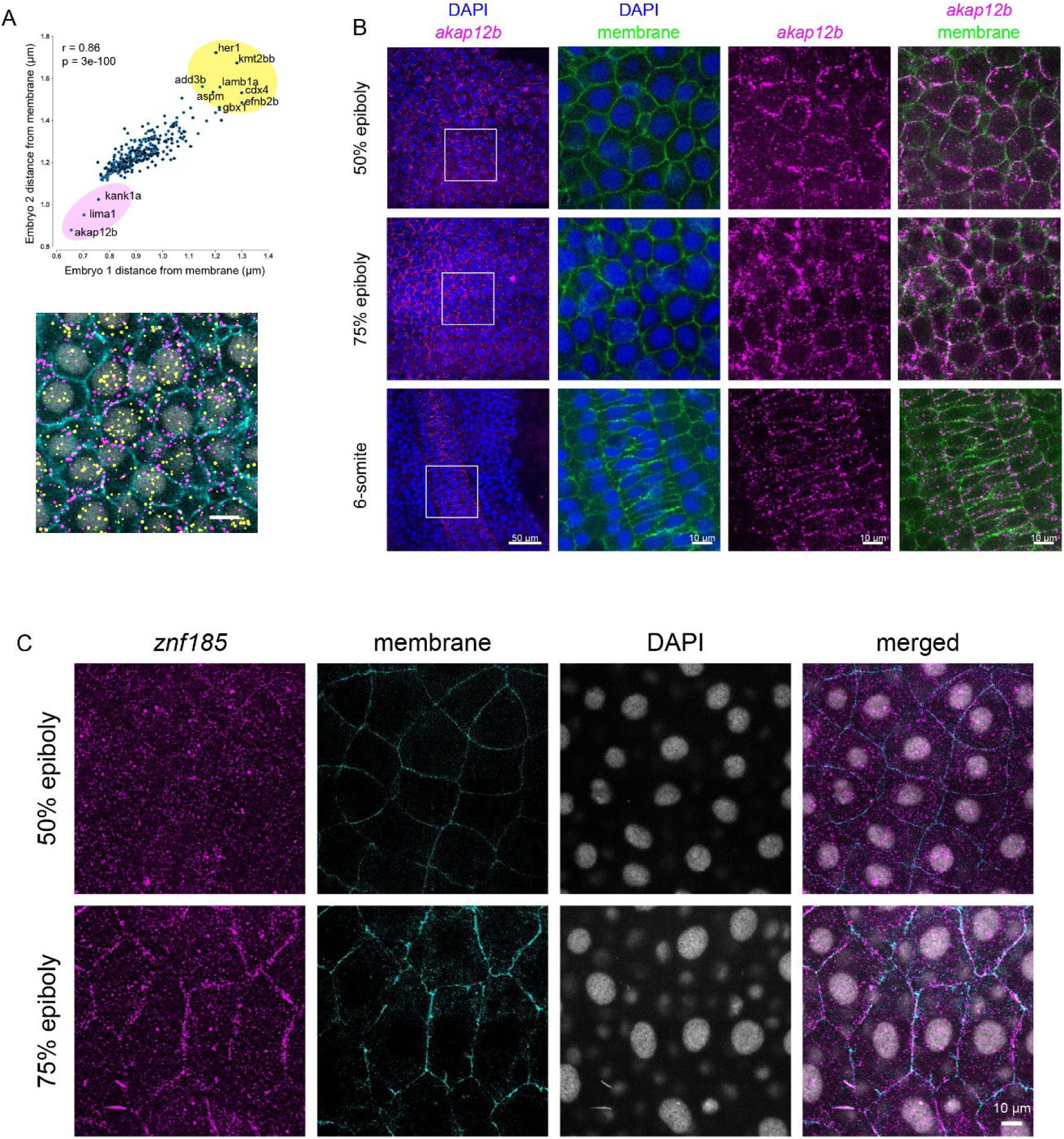
Subcellular localization patterns of weMERFISH data. **(A)** The distance of transcripts from the cell membrane is a conserved feature between embryos, with a high correlation (r = 0.86, p = 3e-100). Each dot in the scatter plot represents the mean distance of a specific gene’s transcripts from the membrane, with error bars showing the standard error of the mean (SEM). Transcripts close to the membrane are marked in magenta, while those further away are in yellow. The accompanying cell image visually confirms these observations. **(B)** *akap12b* transcripts consistently localize to the cell membrane across three developmental stages (50% epiboly, 75% epiboly, and 6-somite). Images display DAPI (nuclei, blue), membrane staining (green), and *akap12b* transcripts (magenta), highlighting their membrane association. **(C)** *znf185* expression in enveloping layer (EVL) cells shows dynamic localization: at 50% epiboly, *znf185* is diffusely distributed, while by 75% epiboly, it becomes concentrated at cell junctions. Sequential images illustrate this transition, with *znf185* in magenta, membrane staining in cyan, and DAPI in gray.

**Fig. S8.**
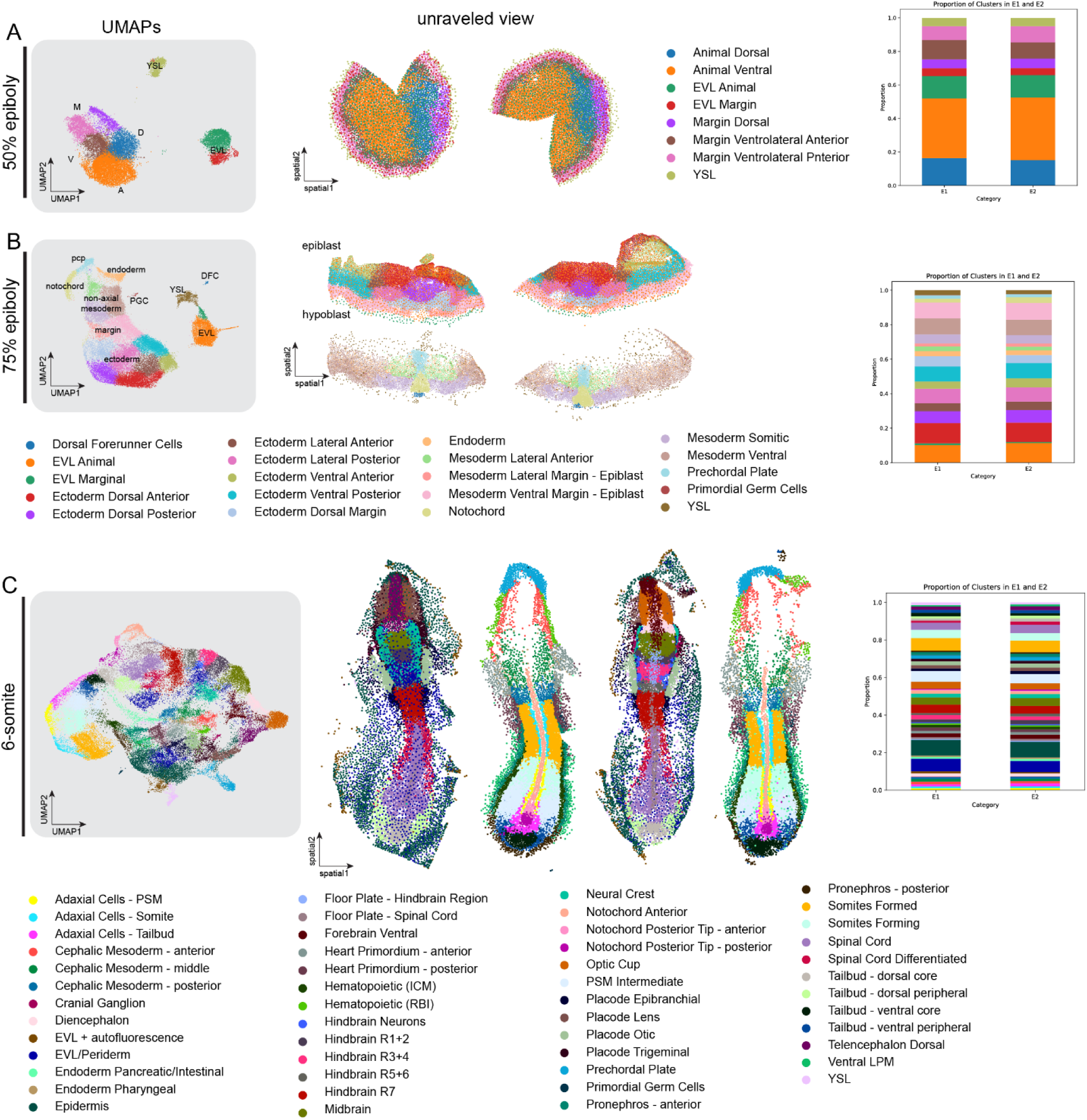
Cell type annotation across replicate embryos. This figure illustrates the cell type annotations for two zebrafish embryos at three key developmental stages: 50% epiboly **(A)**, 75% epiboly **(B)**, and 6-somite **(C)**, using UMAP projections and unraveled views, along with bar plots showing the proportion of cell types in each embryo. This figure emphasizes the detailed annotation of cell types across different stages of zebrafish development and highlights the reproducibility of cell type identification between embryos.

**Fig. S9.**
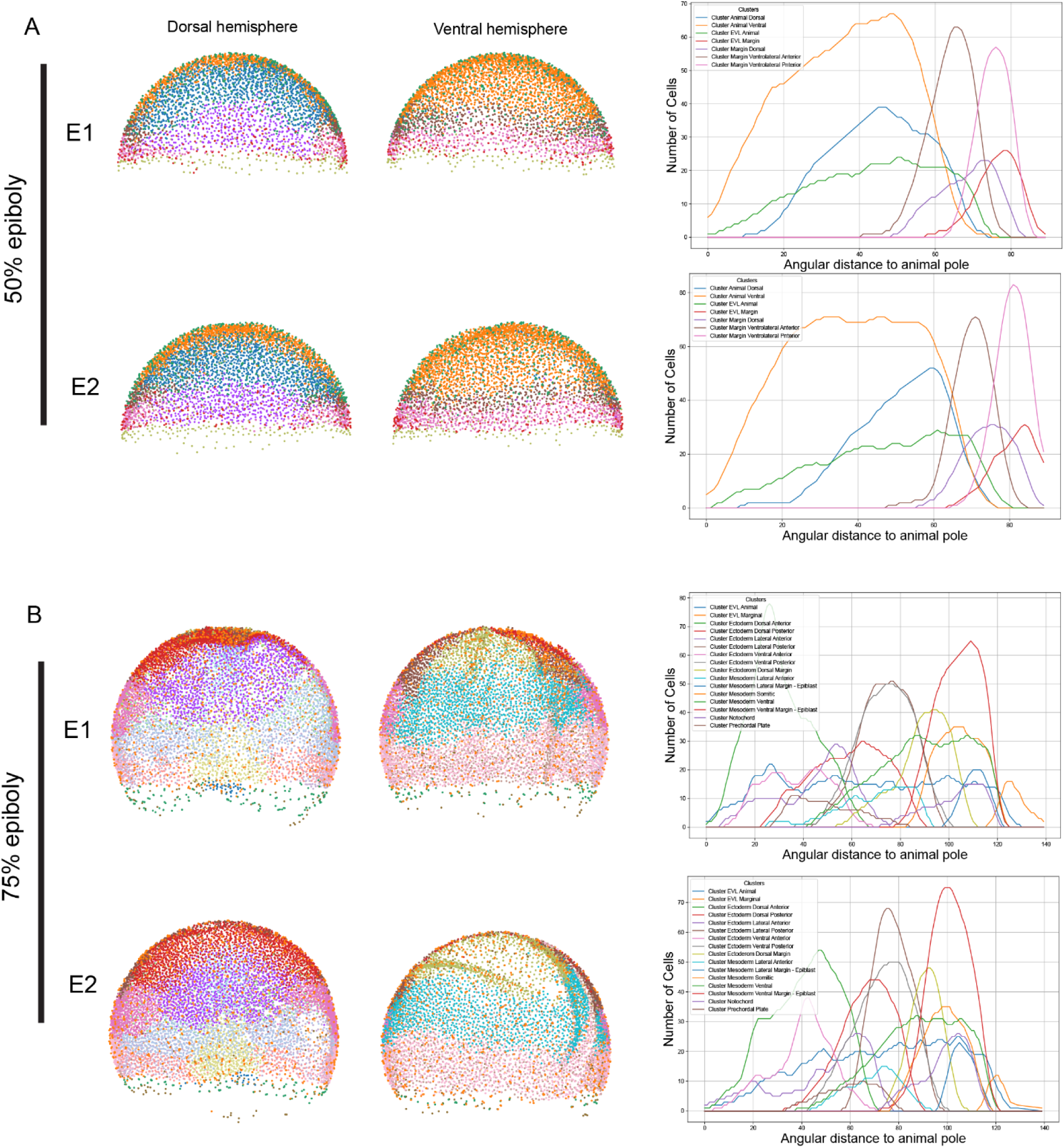
Measurement of location stereotypy across replicate embryos. (A) 50% epiboly: The left panels display the dorsal and ventral hemispheres of embryos E1 and E2, with cells color-coded based on their tissue type. The right panels show plots of the number of cells as a function of their angular distance from the animal pole for different clusters, highlighting the spatial distribution of each cell type in both embryos. **(B)** Same statistics in (A) but for 75% epiboly stage. The plots illustrate how different cell types maintain consistent spatial distributions between the two replicate embryos.

**Fig. S10.**
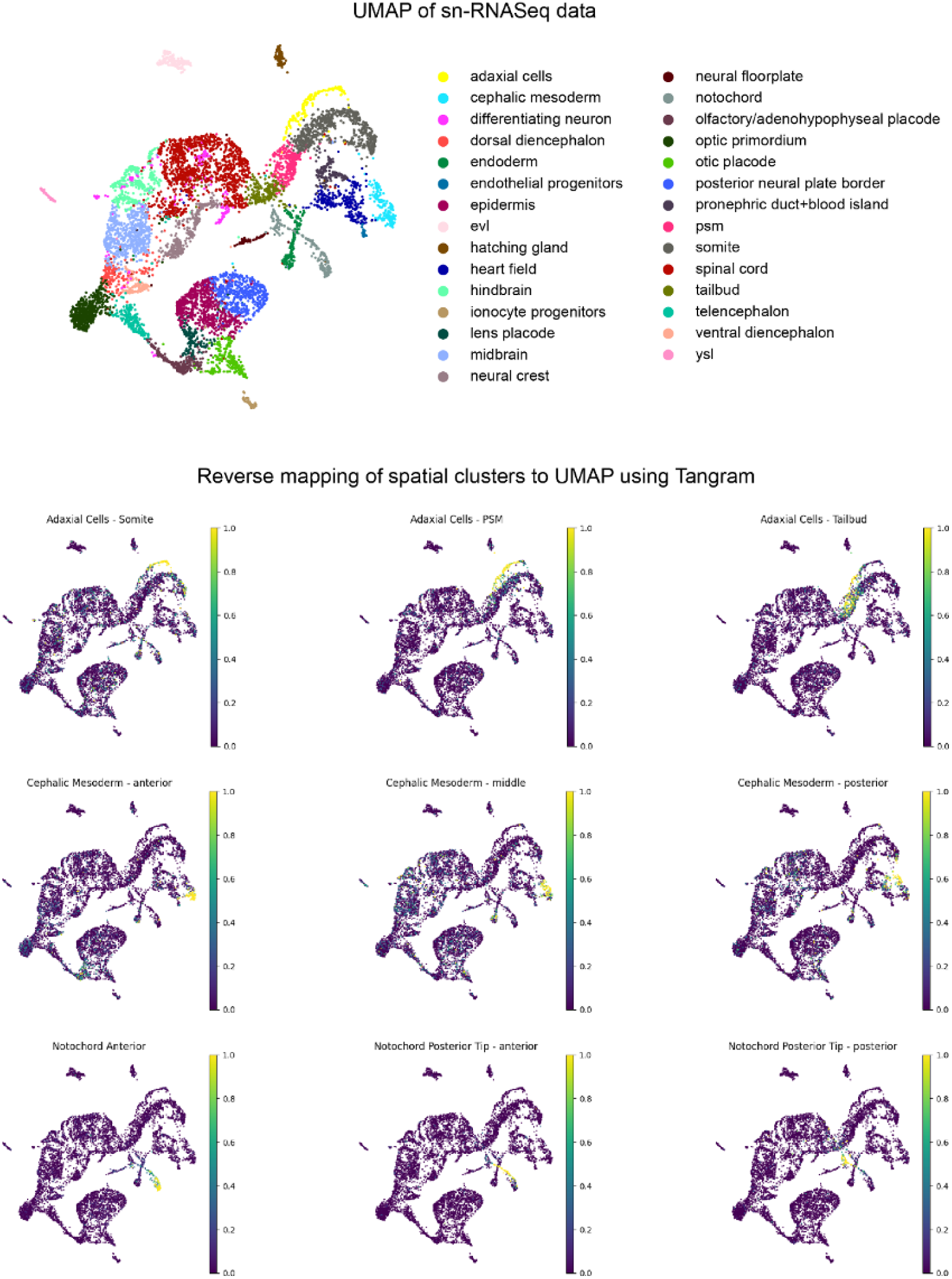
Mapping of spatial subclusters to the UMAP of snRNA-seq data at 6-somite stage. (Top Panel): UMAP of snRNA-seq data: The UMAP shows clustering of various cell types identified in the snRNA-seq data. Each cell type is color-coded as indicated in the legend. (Bottom Panels): Reverse mapping of spatial clusters. The lower panels show the reverse mapping of specific spatial clusters (e.g., adaxial cells in the somite, cephalic mesoderm at various anterior-posterior positions, and notochord regions) onto the UMAP. Each panel visualizes the probability or confidence of cells in the spatial clusters being assigned to corresponding cell types in the UMAP. The color gradient from purple (low probability) to yellow (high probability) indicates the strength of the mapping.

**Fig. S11.**
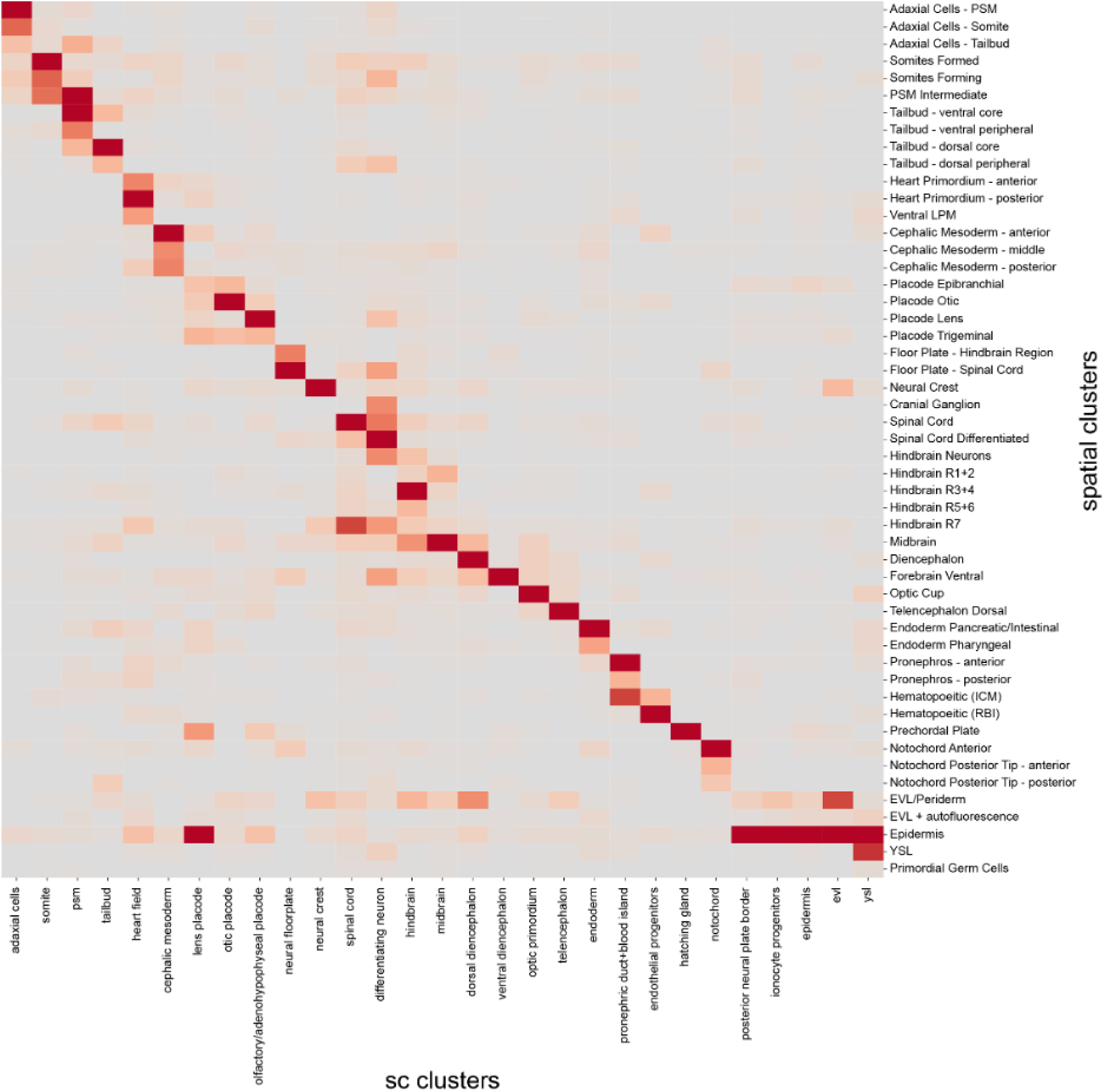
Correspondence of snRNAseq and weMERFISH clusters. This figure presents a heatmap showing the correspondence between snRNA-seq clusters and spatial clusters identified through weMERFISH in 6-somite stage zebrafish embryos. The x-axis represents cell type clusters identified from snRNA-seq data. The y-axis represents spatial clusters identified from weMERFISH data. Each cell in the heatmap reflects the degree of correspondence between a specific snRNA-seq cluster and a spatial cluster, with darker shades of red indicating a higher correspondence.

**Fig. S12.**
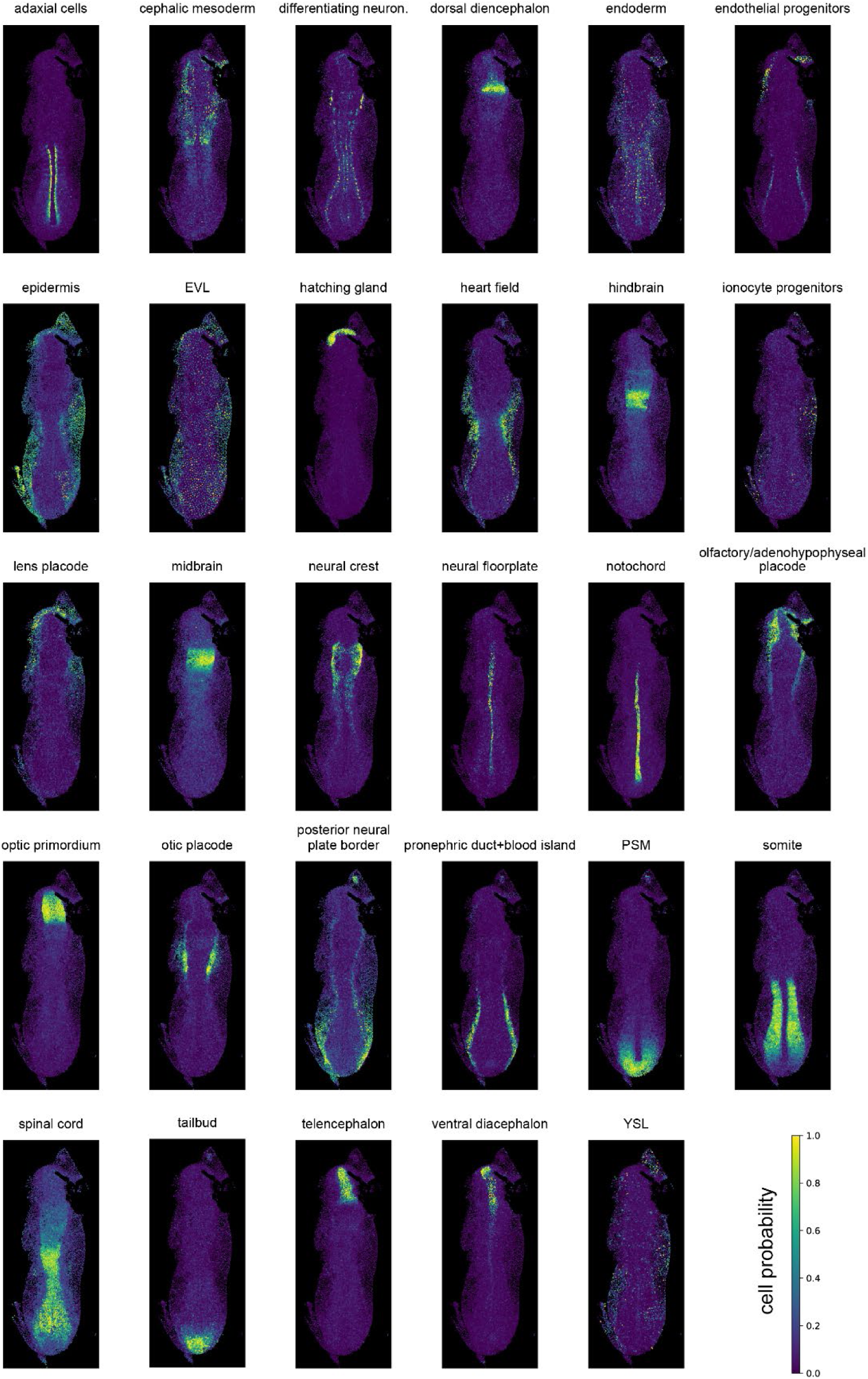
Spatial distribution of snRNASeq cell types. Each panel shows the spatial localization of a specific cell type, projected onto a 2D view of the zebrafish embryo. The color gradient from purple (low probability) to yellow (high probability) represents the likelihood or probability of a cell at a given location in the embryo belonging to the specific cell type shown.

**Fig. S13.**
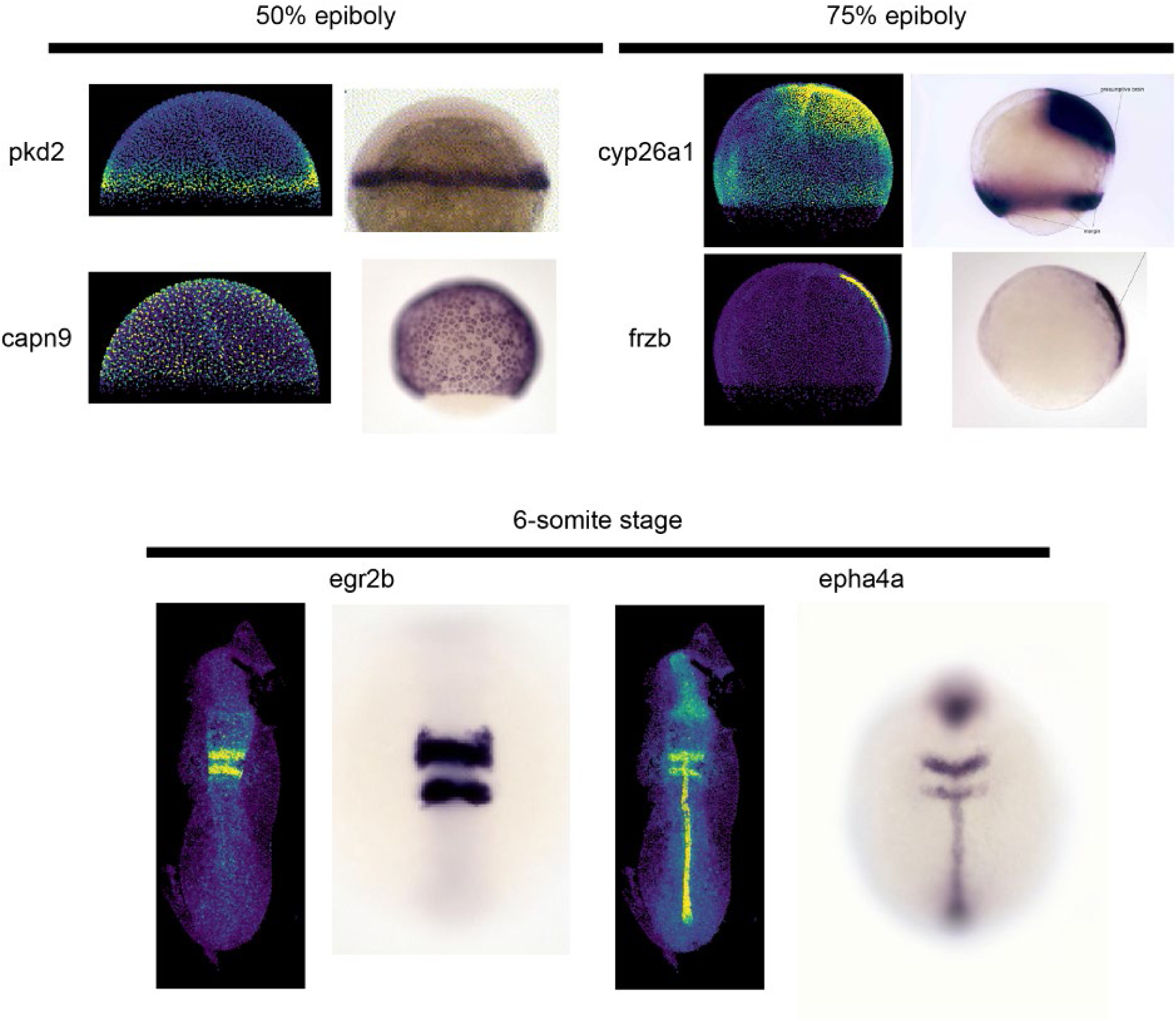
Comparison of imputed gene patterns with published data. Comparison between imputed gene expression patterns from weMERFISH data and published *in situ* hybridization results at three zebrafish developmental stages: 50% epiboly, 75% epiboly, and 6-somite stage. At 50% epiboly, the expression patterns of *pkd2* and *capn9* are compared, showing consistency between the imputed and published data. At 75% epiboly, *cyp26a1* and *frzb* show similar spatial expression patterns in both the imputed and *in situ* data. At the 6-somite stage, *egr2b* and *epha4a* expression patterns in the imputed data closely match the published *in situ* results, demonstrating the accuracy of the imputed spatial gene expression patterns.

**Fig. S14.**
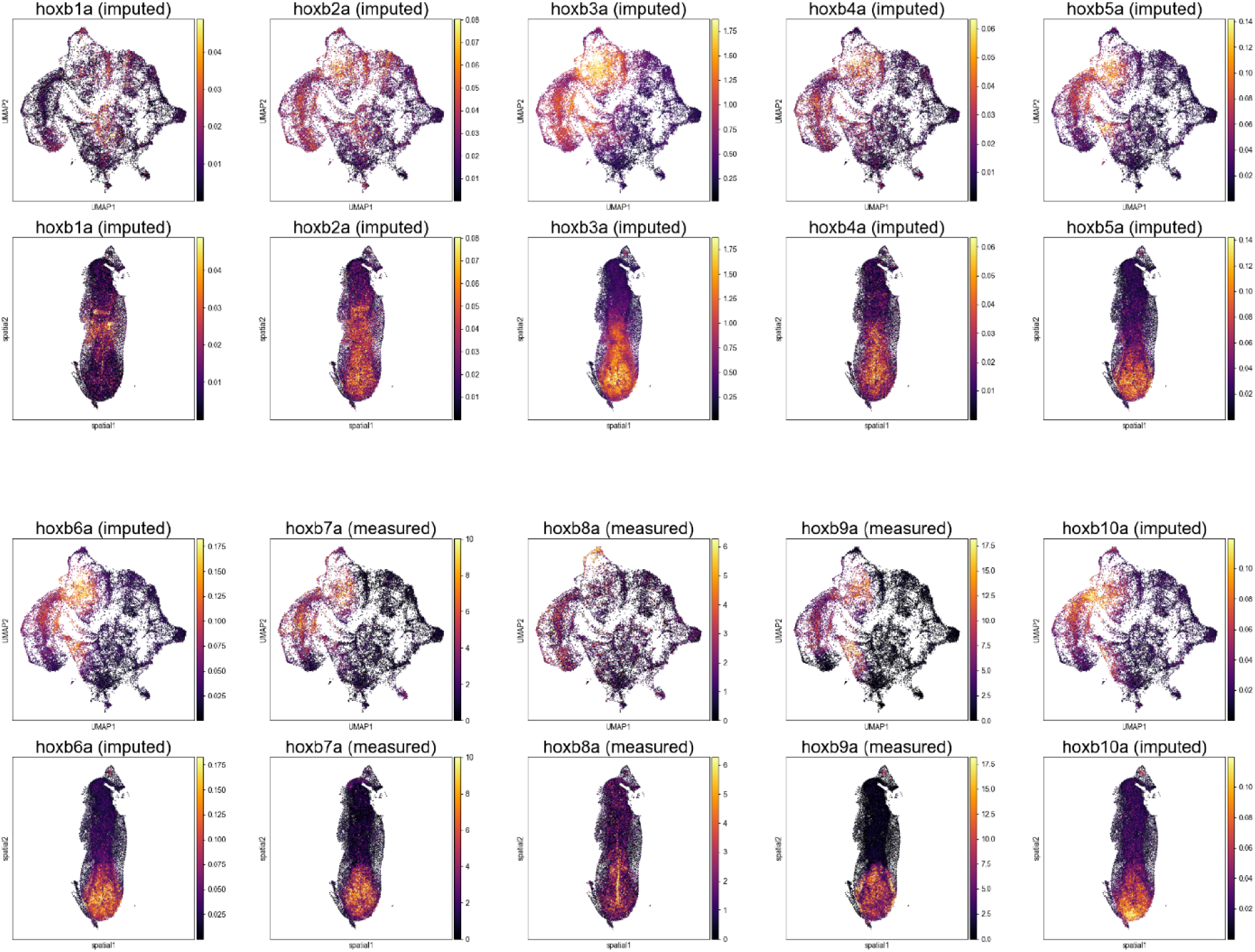
Expression patterns of genes in the Hoxba clusters. The expression patterns of genes in the Hoxba cluster, both imputed and measured, across zebrafish embryos. The top two rows show UMAP and spatial visualizations for the imputed expression of *hoxb1a*, *hoxb2a*, *hoxb3a*, *hoxb4a*, and *hoxb5a*. The bottom two rows display similar visualizations for *hoxb6a*, and *hoxb7a*, *hoxb8a*, and *hoxb9a*, and *hoxb10a*. The color gradient represents the expression level, with yellow indicating higher expression and purple indicating lower expression. These visualizations highlight the spatial and UMAP-based distribution of Hoxba gene expressions in the developing zebrafish embryos.

**Fig. S15.**
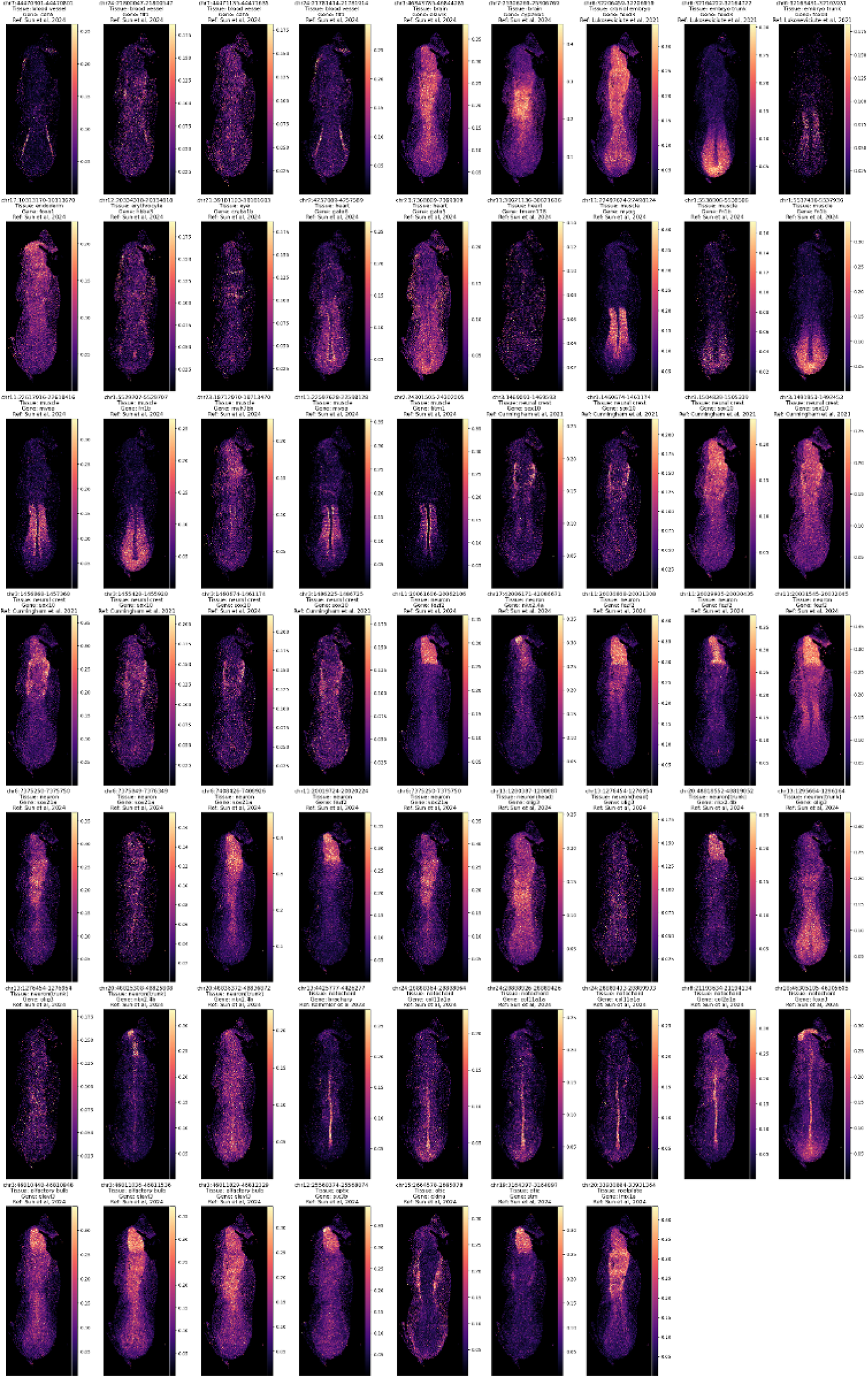
Comparison of imputed ATAC peaks with existing literature. Each panel shows the spatial distribution of chromatin accessibility peaks across different regions of the zebrafish embryo. The imputed peaks are represented with a color gradient, where yellow indicates higher chromatin accessibility and purple indicates lower accessibility.

**Fig. S16.**
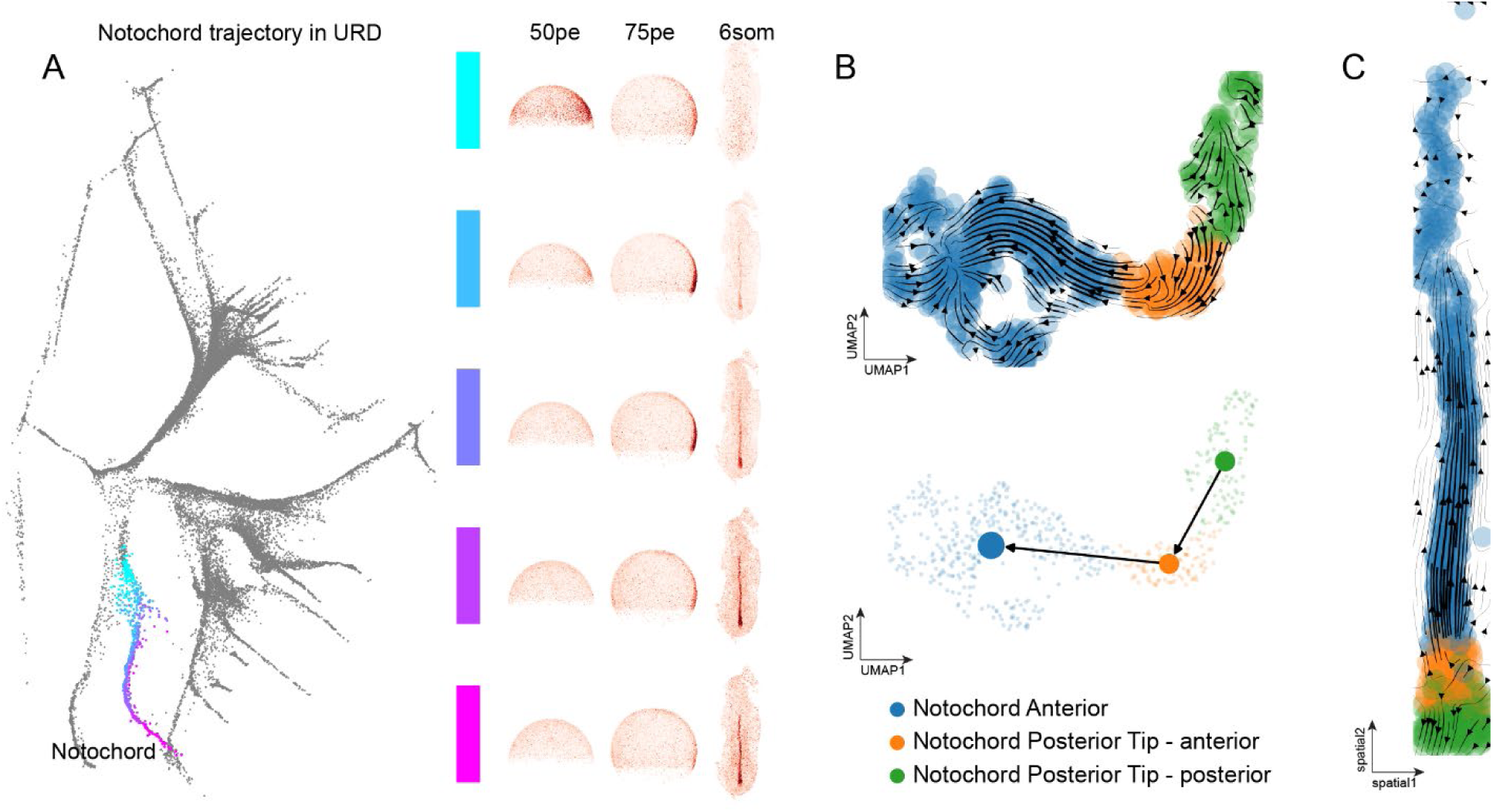
Temporal inference of the trajectory of notochord (A) Notochord pseudotime trajectory mapped to space, showing somite lineages originating at the dorsal margin at 50% epiboly, internalized mesoderm at 75% epiboly, then progress posterior to anteriorly at 6-somtie stage. (B) Schematic illustration of RNA-velocity using nuclear and cytosolic count. (C) RNA velocity inferred for the notochord trajectories using scVelo (upper left) an scVelo + PAGA (lower left). The velocity streamed mapped to space is shown on the right.

**Fig. S17.**
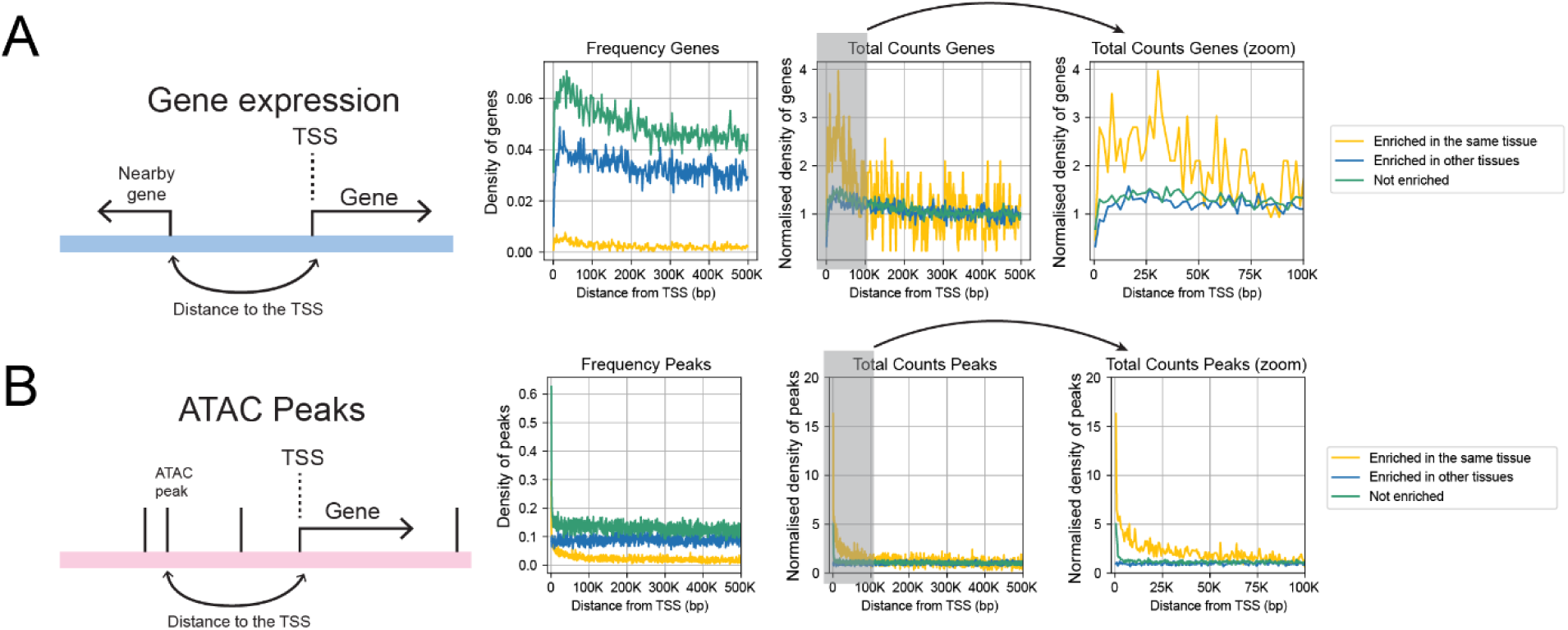
Tissue-specific gene expression and chromatin accessibility patterns **(A)** Gene expression: The enrichment of genes uniquely expressed in a specific tissue is plotted against their genomic distance from the TSS. The line plots show the density and normalized counts of genes enriched in the same tissue (yellow), in other tissues (blue), and not enriched (green). A zoomed-in view of the first 100 kb from the TSS is also provided. **(B)** ATAC peaks: The enrichment of ATAC peaks relative to the TSS is plotted similarly. The line plots display the density and normalized counts of peaks associated with genes enriched in the same tissue (yellow), in other tissues (blue), and not enriched (green), with a zoomed-in view of the first 100 kb.

**Fig. S18.**
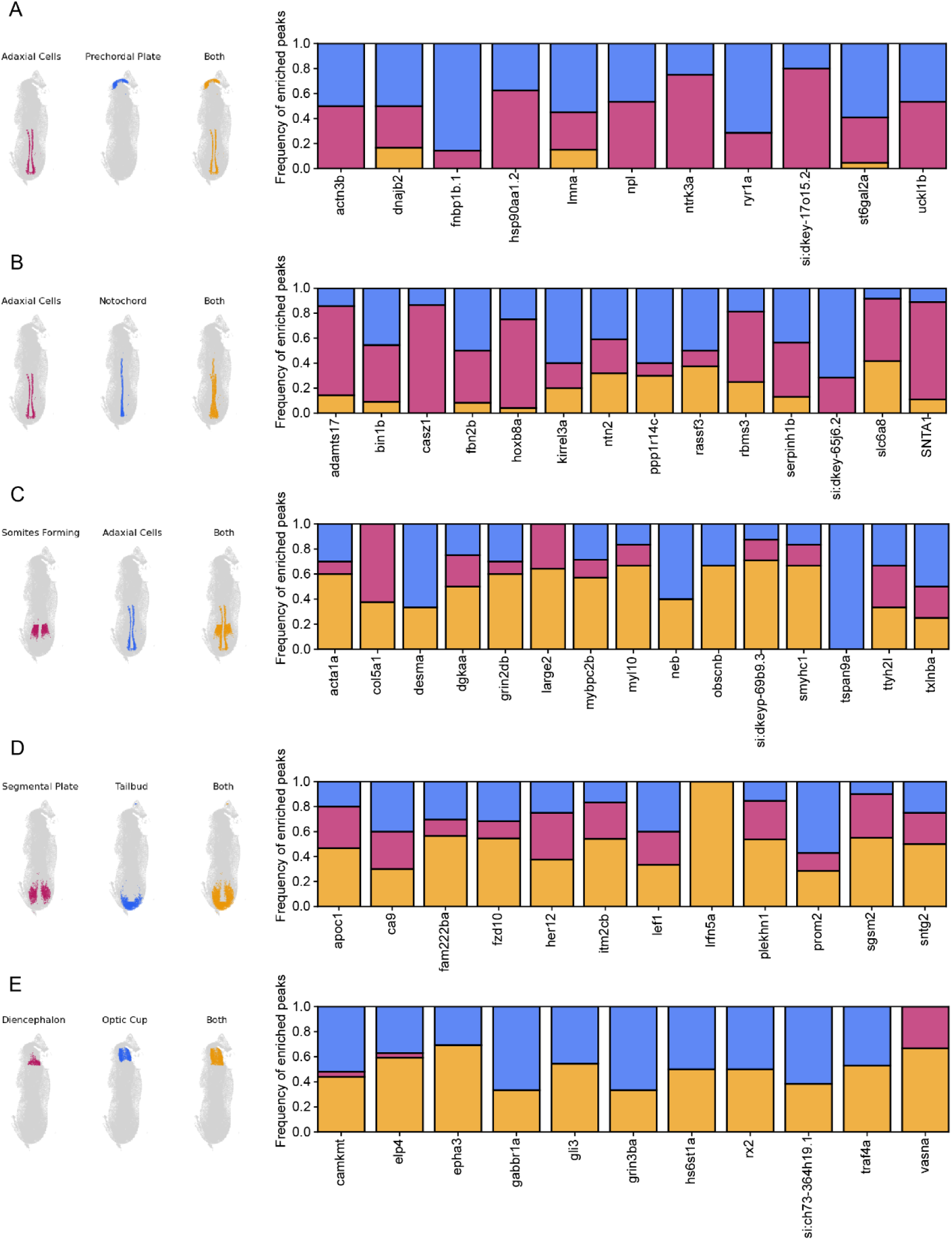
Comparative analysis of regulatory logic of chromatin accessibility in dual-tissue genes. Left: Schematics showing the regions where chromatin is accessible in each pair of tissues. Right: Statistical summary of chromatin accessibility patterns for all genes uniquely expressed in the two tissues. **(A)** Adaxial cells and prechordal plate. **(B)** Adaxial cells and notochord. **(C)** Somites forming and adaxial cells. **(D)** Segmental plate and tailbud. **(E)** Diencephalon and optic cup.

**Fig. S19.**
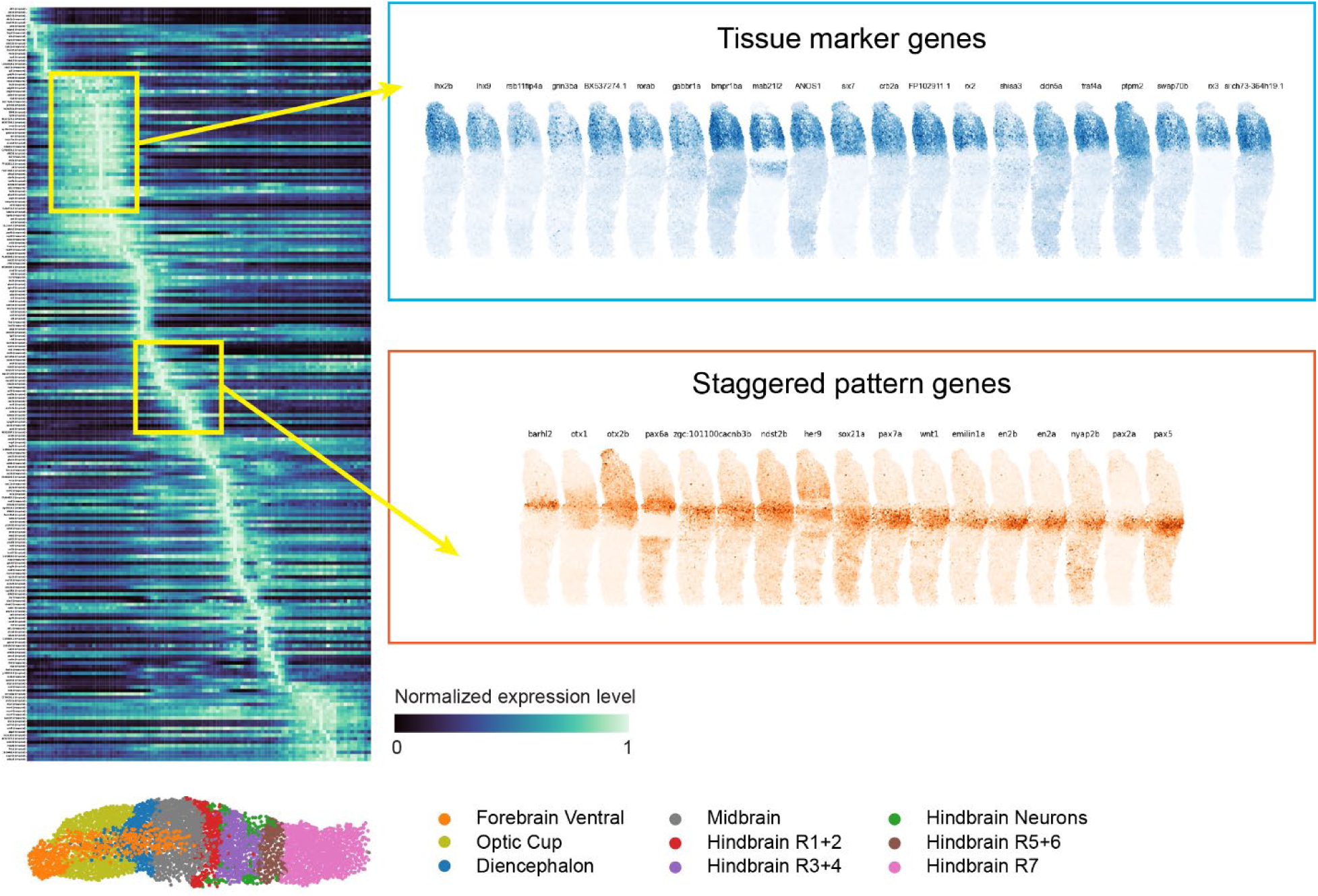
Gene expression in the brain region of 6-somite stage embryo. Heatmap showing two categories of genes: those with sharp borders marking distinct tissues, and those with staggered pattern across anterior-posterior locations

**Fig. S20.**
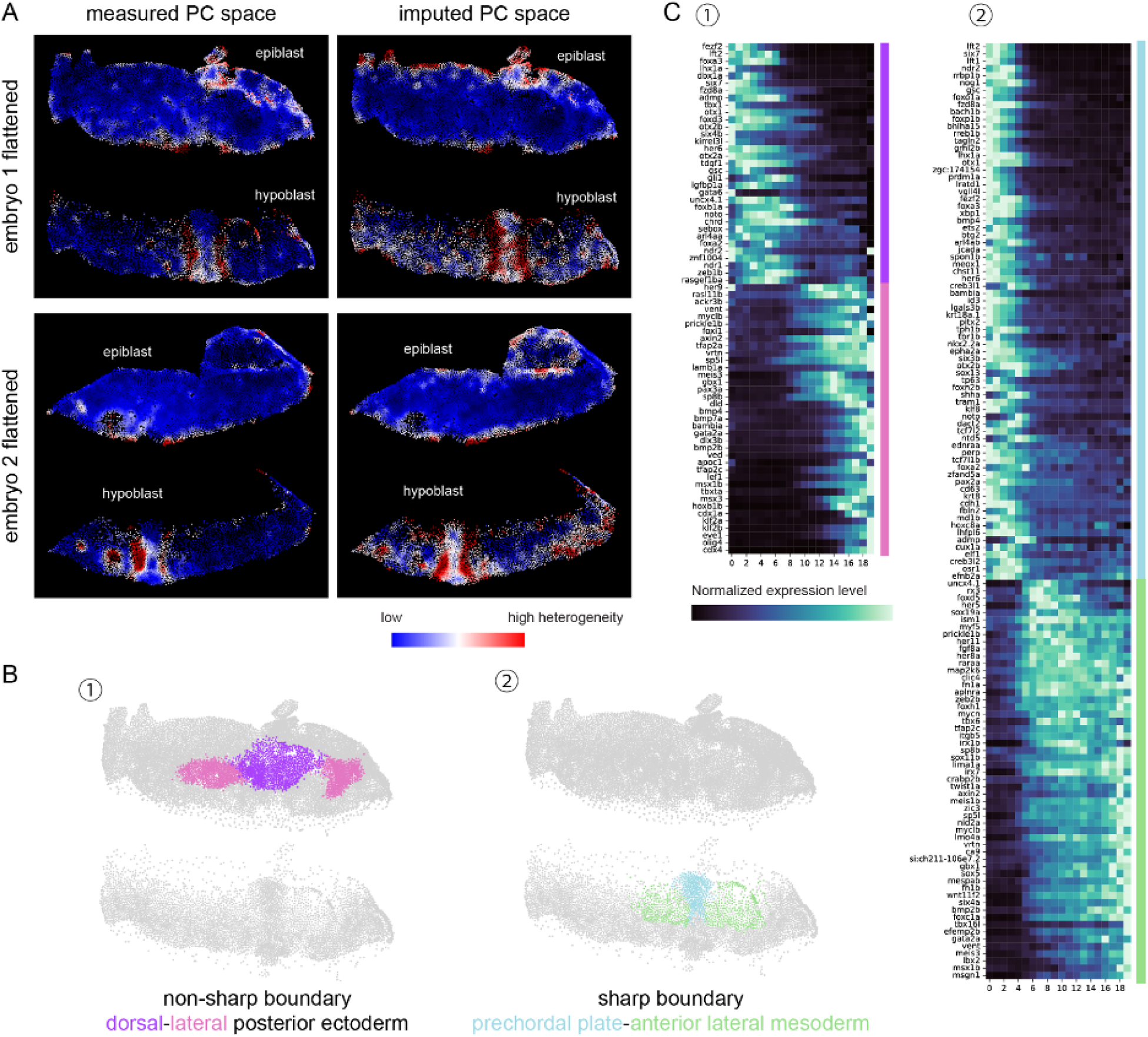
Quantification of boundary sharpness and gene expression across sharp and non-sharp boundary at 75% epiboly stage. **(A)** quantification of the boundary sharpness in flattened view of two embryos. The notochord-somite border emerges as the sharpest boundary in both embryos in both measured and imputed gene expression space. **(B)** Illustration of another non-sharp and sharp boundary at 75% epiboly stage. **(C)** Gene expression across the non-sharp boundary at dorsal-lateral posterior ectoderm border and sharp boundary at the prechordal plate-anterior lateral mesoderm border

**Fig. S21.**
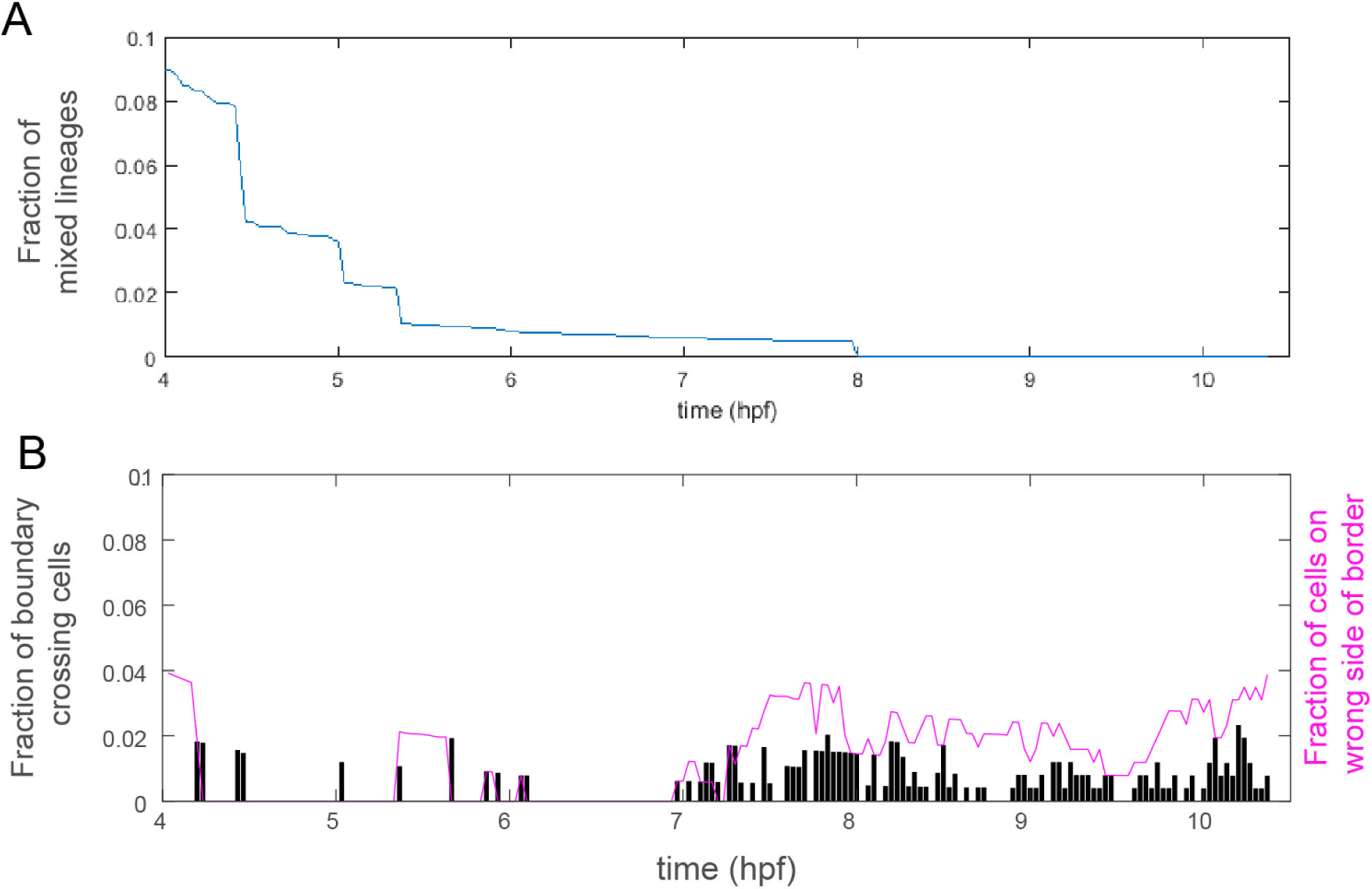
Characterization of lineage and cell behaviors during notochord-somite boundary formation (A) Fraction of mixed lineages during notochord-somite boundary formation (B) Quantification of cell behaviors as in number of cells that cross the border of straight line fitted between the population of cells of the notochord and somite fates. The cells on wrong side of the border by the end of time lapse imaging are likely artifact that the border is not a straight line.

**Movie S1.**

Light-sheet imaging of notochord-somite boundary formation

**Data S1. (separate file)**

Literature evidence of 6-somite stage cell type markers

**Data S2. (separate file)**

Genes in the 1000-gene library

**Data S3. (separate file)**

Sequences of the amplification and readout probes

**Data S4. (separate file)**

List of cross-reference to published enhancers

